# p53 activity licenses transcriptional regulation by YAP/TAZ-TEAD to shape expression landscapes during tumorigenesis

**DOI:** 10.1101/2025.09.29.679387

**Authors:** Cole Martin, William B. Sullivan, Scott Bang, Joseph R. Gomes, A. Cole Edwards, Channing J. Der, Cheng Fan, Yusha Liu, John P. Morris

## Abstract

The tumor suppressor p53 is the most frequently disabled gene in cancer, where its inactivation promotes the transition between transcriptionally distinct pre-malignant and malignant disease. How p53 action shapes these stage-dependent expression landscapes is poorly understood. Using a mouse model where p53 function is restored in advanced pancreatic cancer, we demonstrate that p53 dependent gene expression is determined by regulation of YAP/TAZ-TEAD transcriptional complexes at multiple levels of molecular control. Restoration of p53 activity in pancreatic cancer cells expands transcriptional regulation by reorganizing TEAD binding at newly accessible enhancer landscapes, resulting in distinct TEAD-driven gene regulatory networks when p53 is inactivated versus engaged. Furthermore, p53 potentiates expression of newly licensed TEAD targets by increasing TAZ levels in response to the actin remodeling associated with cell cycle arrest and senescence. Thus, p53 activity controls the qualitative and quantitative output of YAP/TAZ-TEAD, including secretory programs involved in tumor-immune communication and remodeling of the microenvironment. Our work nominates regulation of YAP/TAZ-TEAD via p53 as a mechanism that determines stage dependent transcriptional landscapes during stepwise tumorigenesis.

## INTRODUCTION

Inactivation of the tumor suppressor p53 is the most common event in cancer, representing a path to malignancy in more than half of all tumors^1,2^. Loss of p53 acts as a genetic inflection point during cancer progression, collaborating with driver mutations that initiate pre-malignant disease to unleash the progression of indolent precursors to frank malignancy^2–5^. Inactivation of p53 triggers biological and transcriptional diversification^6,7^ during malignant initiation, implicating a role for p53 function in determining how cells establish distinct expression states during tumorigenesis. However, the molecular mechanisms connecting p53 activity to the transcriptional heterogeneity that characterizes these phenotypic transitions during the pre-malignant to malignant switch remain poorly understood.

Pancreatic ductal adenocarcinoma (PDAC) is a lethal cancer where p53 is the primary, genetic barrier constraining malignant progression^3,5,8,9^. The genetic landscape of PDAC is dominated by nearly universal, initiating mutations in KRAS that are followed by inactivating mutations in TP53 in ∼75% of cases^10^. Loss of p53 function is strongly associated with progression to transcriptionally heterogeneous frank disease^11,12^. The role of p53 in constraining malignant progression is highly conserved, as sporadic, biallelic inactivation of p53 in mouse PDAC models driven by mutant Kras is a nearly obligate event in the pre-malignant to malignant switch^3,5^. p53 inactivation results in the development of heterogeneous, epigenetically diverse lineages, characterized by altered accessibility of transcription factors that play context dependent roles in pre-malignant and malignant cells^13,14^.

We have previously demonstrated that p53 function actively constrains malignant progression in Kras mutant PDAC, where restoration of p53 in malignant cells triggers global remodeling of chromatin accessibility, enforces pre-malignant gene expression, and induces features of pre-malignant differentiation^15^. Here we demonstrate that p53 establishes these stage-dependent gene expression programs in part by regulating the function of YAP/TAZ-TEAD transcriptional complexes, which are critical determinants of both PDAC initiation and progression^16,17^, at multiple levels of molecular control. TEAD regulatory networks are qualitatively shaped via reorganization of TEAD genomic binding at chromatin landscapes remodeled in response to p53. Furthermore, p53 determines the magnitude of TEAD target gene expression via accumulation of its co-activator TAZ in response to the actin reorganization accompanying cycle arrest and cellular senescence. p53-dependent TEAD regulation thus coordinates distinct regulomes that reflect differences in pre-malignant and malignant gene expression during PDAC progression. Among these are secretory programs that are also selectively regulated by TEAD in response to senescence induced by KRAS/MAPK pathway inhibitors. Our work highlights the coordination of TEAD transcriptional regulation in response to p53 and its effects on cell fate as a novel route by which p53 function can establish context-dependent gene regulation and direct stage-dependent biology in tumorigenesis.

## RESULTS

### p53 activity remodels chromatin to diversify YAP/TAZ-TEAD enhancer binding in pancreatic cancer cells

The malignant switch unleashed by p53 inactivation in PDAC development is characterized by distinct chromatin landscapes and stage-dependent accessibility of transcription factors (TFs) that direct neoplastic fate during PDAC progression^13,14,18^. To ask if p53 plays an active role in establishing these stage-dependent transcriptional landscapes, we analyzed chromatin accessibility via ATAC-SEQ in response to restoration of wildtype p53 function in primary KP^sh^ pancreatic cancer cells. In this model, p53 is silenced during pancreatic cancer development via a doxycycline (dox)-induced shRNA expressed specifically in the pancreatic epithelium upon Cre-mediated activation of latent, conditional alleles permitting expression of mutant Kras^G12D^ and a reverse tetracycline-controlled transactivator (rtTA) (KP^sh^: *Pdx1-Cre; LSL-Kras^G12D^; TRE-shp53; Rosa26-LSL-rtTA-IRES-mKate2*^15,19^). Dox-fed mice display accelerated malignant progression that mirrors carcinogenesis driven by irreversible, conditional inactivation of p53 in Kras mutant pancreatic cells and that recapitulates key histopathological features of human disease^15,19^(Suppl. Fig.1A). Dox withdrawal from primary KP^sh^ PDAC cell lines results in robust accumulation of transcriptionally active p53, as demonstrated by potent induction of well characterized p53 direct target genes such as Cdkn1a/p21, and induction of p53 triggered changes in cell fate including cell cycle arrest corresponding with markers of cellular senescence (e.g., senescence-associated β−galactosidase, SA-β−Gal, Suppl. Fig.1B-D).

Sustained p53 restoration in KP^sh^ cells resulted in widespread remodeling of global chromatin landscapes with increased accessibility at ∼24% (21,676/91,191) and reduced accessibility at ∼28% (25,698/91191) of identified peaks, respectively (fold change>1.5, FDR<0.05) (Fig. 1A). Motif enrichment analysis of loci opened in response to p53 restoration revealed significant enrichment of response elements recognized by TFs implicated in both pre-malignant and malignant states in both mouse and human PDAC^13^, e.g., AP-1, TEAD, FOXA1, and BACH (Fig.1B). To ask if p53 triggered chromatin landscapes are reflective of regulatory networks observed in the pre-malignant to malignant switch defined by loss of p53, we compared motif enrichment patterns following p53 restoration in KP^sh^ cells to those observed in previously published chromatin accessibility mapping of Kras mutant, p53 wildtype pre-malignant cells versus Kras mutant, p53 inactivated PDAC^13^. Implicating a direct role for p53 activity in determining TF regulatory landscapes during neoplastic progression driven by mutant Kras, we observed remarkable consistency in motifs enriched in loci opened in association with wildtype p53 in both models, with AP-1, TEAD, NR-F2, and BACH amongst the most enriched in both systems and >70% overlap in significantly enriched motifs (Fig. 1B-D).

**Figure 1.**
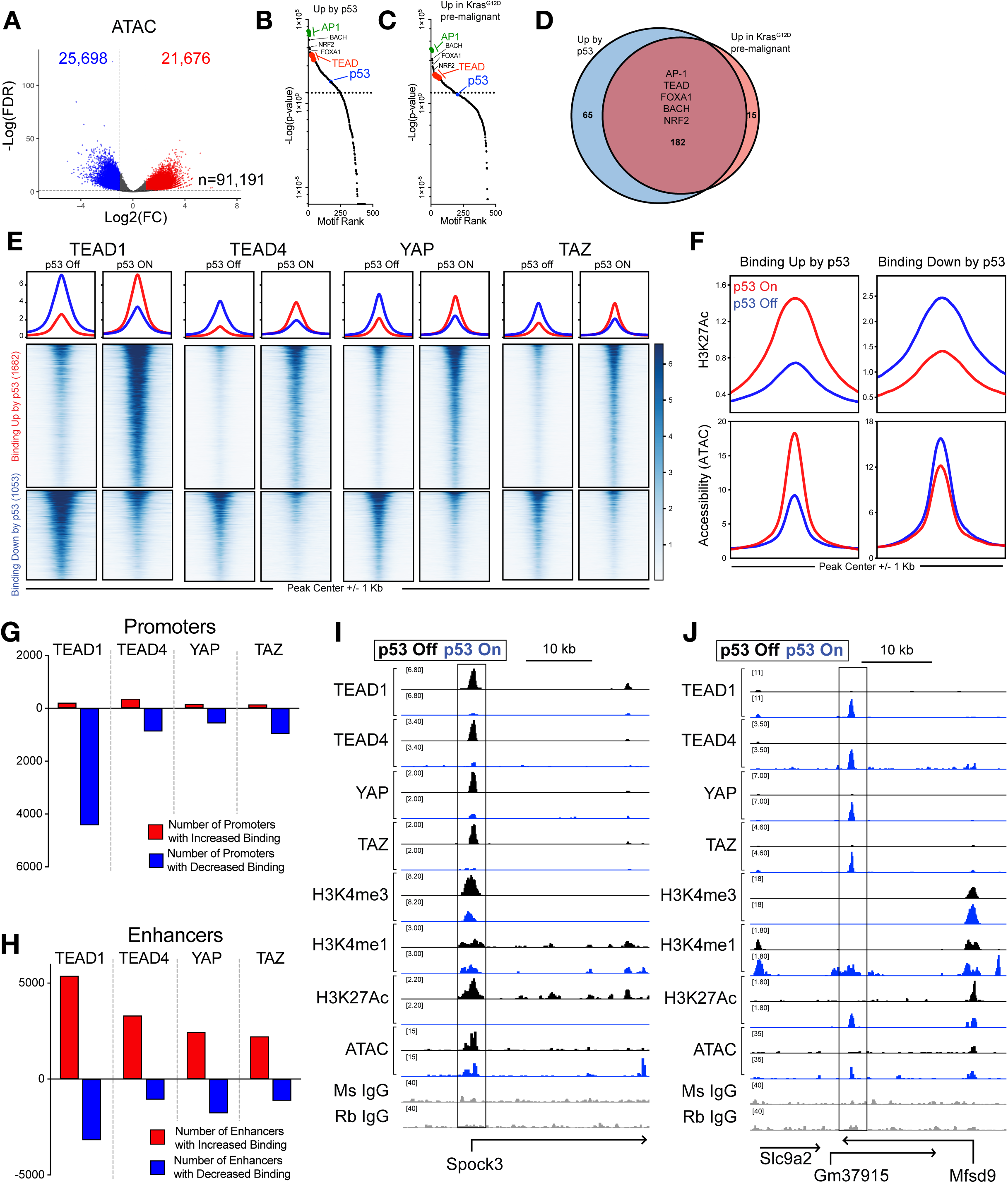
p53 reshapes the YAP/TAZ-TEAD genomic binding landscape. **(A)** Volcano plot of differentially accessible peaks (EdgeR) in KP^sh^1 cells on dox or 8 days off dox (p53 on). Red and blue dots denote peaks with significantly increased and decreased openness respectively (FC>1.5, FDR<0.05). **(B-C)** Rank ordered plot of significantly enriched motifs (FDR<0.05) in loci with opened accessibility (FC>1.5, FDR<0.05) **(B)** upon 8 days of p53 restoration in KP^sh^1 or **(C)** in pre-malignant p53 wild type pancreatic metaplasia over malignant p53 inactivated mouse PDAC. **(D)** Venn diagram of overlapping significantly enriched motifs from (B) and (C). **(E)** Normalized peak densities of shared peaks identified in CUT&RUN for TEAD1, TEAD4, YAP, and TAZ with differential binding upon restoring p53 detected in all four proteins (FC>1.5, FDR<0.05). Top row showing profile plots of binding magnitude with red and blue lines indicating peaks with shared increased and decreased binding respectively. Middle row and bottom row showing heatmaps of peaks with shared increased and decreased binding upon restoring p53 respectively. **(F)** Profile plots of normalized H3K27Ac abundance (CUT&RUN, top) and accessibility (ATAC-seq, bottom) over loci with shared increased (left) or decreased (right) binding magnitude of TEAD1, TEAD4, YAP, and TAZ upon restoring p53 (FC>1.5, FDR<0.05). Blue and red lines indicate signal on dox (p53 off) or 6 days off dox (p53 on) respectively. **(G)** Graph showing the number of promoter peaks (assigned as overlapping a H3K4me3 peak and within 1 Kb of a known TSS) with differential binding upon p53 restoration (FC>1.5, FDR<0.05). **(F)** Graph showing the number of enhancer peaks (assigned as overlapping a H3K4me1 peak and not within 1 Kb of a known TSS) with differential binding upon p53 restoration (FC>1.5, FDR<0.05). **(I-J)** Tracks showing normalized CUT&RUN for TEAD1, TEAD4, YAP, and TAZ on dox (p53 off, black), 6 days off dox (p53 on, blue), or isotype matched IgG (grey) at an example of a differentially bound **(I)** promoter (Spock3 TSS) or **(J)** enhancer (Mfsd9 intron).

We next asked if the accessibility landscapes observed in response to p53 reflect functional reorganization of TF regulatory networks. TEAD complex occupancy was interrogated due to the important role of the paralogous TEAD coactivators Yes-associated protein (YAP) and transcriptional coactivator with PDZ binding motif (TAZ)^20^ in both the development of Kras driven pre-malignant disease^16,21^ and the progression, maintenance, and therapeutic response of advanced, p53 mutant PDAC^17,22–24^). Therefore, we performed CUT&RUN profiling of TEAD1, TEAD4, YAP, and TAZ in KP^sh^ cells following p53 restoration. Profiling was robust as genomic occupancy was highly overlapping across TEAD complex members, tracked closely with p53 activity, and revealed robust binding at the promoter of the well characterized, direct YAP/TAZ-TEAD target genes such as Cyr61 both on and off dox (Suppl. Fig. 2A-C).

Restoring p53 function produced dramatic changes in genomic binding across all four TEAD complex components, with differential binding identified at 50.7% of TEAD1 peaks, 37.3% of TEAD4 peaks, 30.1% of YAP peaks, and 31.4% of TAZ peaks (fold change>1.5, FDR<0.05) (Suppl. Fig. 2D). TEAD family members share a response element and frequently display redundant binding patterns that are occupied by complexes containing either YAP or TAZ^25^. Consistent with overlapping occupancy in other cell types, we observed strong correlation in differential binding between TEAD1 and TEAD4 (R^2^=0.798, p<1e^-15^) as well as YAP and TAZ (R^2^=0.778, p<1e^-15^) triggered by p53 activity (Suppl. Fig.2E,F), with mutually increased binding of all four TEAD complex members upon p53 restoration at 1,682 peaks and shared decreased binding at 1,053 peaks (fold change>1.5 and FDR<0.05) (Fig. 1E). Importantly, context-dependent binding of YAP/TAZ-TEAD complexes corresponded with gained and lost chromatin accessibility following p53 restoration, at loci that were highly enriched for TEAD motifs (Fig. 1F, Suppl. Fig.2G). These data demonstrate that p53 maintains chromatin states that license distinct patterns of transcription factor binding that are characteristic of the pre-malignant versus malignant switch in pancreatic cancer and specifically acts as a contextual determinant of regulatory networks of YAP/TAZ-TEAD transcriptional complexes.

YAP/TAZ-TEAD complexes predominantly regulate gene expression via interpretation of enhancer landscapes, with a minority of direct targets dependent on promoter engagement^25,26^. Furthermore, enhancer remodeling has emerged as an integral determinant of gene expression programs and cell fate changes triggered by p53^27,28^. Thus, we integrated YAP/TAZ-TEAD binding with detailed profiling of histone modifications associated with enhancers and promoters to ask if remodeling of YAP/TAZ-TEAD complex binding reflects engagement with epigenetic landscapes triggered by p53..

We used CUT&RUN to detect the genomic distribution of histone post-translational modifications that define enhancers and promoters with and without p53 restoration: histone 3 lysine 4 trimethylation (H3K4me3, active promoters), histone 3 lysine 4 monomethylation (H3K4me1, enhancers), and histone 3 lysine 27 acetylation (H3K27Ac, active enhancers/promoters)^29^. H3K4me3, H3K4me1, and H3K27Ac profiling displayed robust, p53 dependent clustering and the expected distribution across genomic features (Suppl. Fig 3.A,B). . p53 restoration in KP^sh^ cells broadly altered the abundance of H3K4me3 (∼29%, 7216/24688 peaks), H3K4me1 (∼17%, 5144/30104 peaks), and H3K27Ac (∼30%, 7345/22821 peaks) genome wide (Suppl. Fig. 3C).

Integrating CUT&RUN profiling for TEAD1, TEAD4, YAP, and TAZ with histone modifications demonstrated that genomic reorganization of TEAD complexes triggered by p53 corresponded with engagement of newly established enhancer landscapes. First, we assigned TEAD1, TEAD4, YAP, and TAZ peaks overlapping H3K4me1 peaks >1 kb from transcriptional start sites (TSSs) as enhancers and YAP/TAZ-TEAD peaks overlapping a H3K4me3 peak <1 kb from TSSs as promoters. A minority of YAP/TAZ-TEAD peaks were called at promoter regions where, remarkably, p53 activity resulted in reduced occupancy of TEAD1, TEAD4, YAP, and TAZ (Fig. 1G, Suppl. Fig.3G). On the other hand, p53 triggered a global expansion of YAP/TAZ-TEAD at enhancers, with TEAD1, TEAD4, YAP, and TAZ consistently exhibiting increased binding magnitude following p53 restoration (Fig. 1H). Thus, p53 triggered context-dependent YAP/TAZ-TEAD gene regulation, with a tendency towards exclusion of YAP/TAZ-TEAD binding near transcriptional start sites of otherwise bound genes (e.g., Spock3, Fig. 1I), and potentiated binding at candidate enhancer loci of others (e.g., Mfsd9, Fig. 1J). Importantly, dynamic YAP/TAZ-TEAD binding in response to p53 activity corresponded with the transcriptional activation state of chromatin as indicated by H3K27Ac abundance (Fig. 1F).

Exceptionally high levels of transcriptional activators and active epigenetic marks including H3K27Ac define “super enhancers” which regulate key cell-identity genes^30^ and have been shown to be remodeled in p53-triggered cell fates including cellular senescence^16,31^, Thus, we asked if reorganized YAP/TAZ-TEAD binding in response to p53 action also reflected engagement with dynamically remodeled super-enhancers. We identified 2044 super enhancers upon p53 restoration in KP^sh^ cells, by selecting non-promoter H3K27Ac peaks where the H3K27Ac signal exceeded its rank-ordered geometric mean (Suppl. Fig. 3E). p53 resulted in remodeling of a substantial portion of super enhancers, resulting in increased and decreased H3K27Ac levels at 490 and 224 super enhancers, respectively (fold change>1.5, FDR<0.05, Suppl. Fig. 3F). Remarkably, motif enrichment analysis identified TEAD response elements as the most enriched motifs in super enhancers activated in response to p53 restoration (Suppl. Fig. 3G) and accordingly, TEAD1, TEAD4, YAP, and TAZ binding corresponded with super enhancer opening and closing in response to p53 restoration (Suppl. Fig. 3H)). Taken together, these data identify chromatin remodeling triggered by p53 as a contextual determinant of YAP/TAZ-TEAD genomic binding, largely at enhancer regulatory networks.

### p53 licenses distinct YAP/TAZ transcriptional targets in pancreatic cancer cells that reflect pre-malignant and malignant expression programs defined by p53 status in PDAC development

Because p53 activity results in reorganization of YAP/TAZ-TEAD binding, we next asked if p53 and YAP/TAZ cooperatively establish gene expression programs in Kras mutant pancreatic cancer cells. We performed RNA-seq in KP^sh^ cells constitutively expressing shRNAs that silence either Yap or Taz, maintained on dox or 6 days after dox withdrawal. Importantly, these shRNAs resulted in effective Yap and Taz knockdown at the mRNA and protein levels before and after restoring p53, with p53 protein levels unchanged by either Yap or Taz depletion (Suppl. Fig. 4A-D). Yap and Taz knockdown effectively reduced TEAD activity in p53 off and on states as demonstrated by reduced expression of the YAP/TAZ-TEAD direct targets Cyr61 and Aldh1a (Suppl. Fig. 4E).

Consistent with a well-established role for Yap and Taz dependent transcriptional regulation during pancreatic cancer progression^16,17^, Yap or Taz knockdown altered gene expression in both p53 silenced and active states (Fig. 2A), and regulated similar sets of genes in both contexts, consistent with their largely overlapping genomic binding landscapes (Suppl. Fig. 4F). However, PCA analysis revealed a much stronger effect of Yap and Taz knockdown on global gene expression following p53 restoration (Fig. 2A). Accordingly, we identified 4.5-fold more differentially expressed genes (DEGs) after restoring p53 in cells expressing shTaz (95 p53 off vs 432 p53 on, Fig. 2B) and 5.3-fold more DEGs in cells expressing shYap (60 p53 off vs 843 p53 on) (fold change>1.5, FDR<0.05, Fig. 2C). Consistent with expanded regulation by YAP/TAZ in p53 on cells, gene set enrichment analysis (GSEA) demonstrated more robust target gene depletion in shYap and shTaz cells after restoring p53 across multiple YAP/TAZ-TEAD target gene signatures (Fig. 2D). Thus, p53 acts to expand and diversify YAP/TAZ gene regulatory networks in pancreatic cells, licensing regulation of distinct sets of YAP/TAZ target genes.

**Figure 2.**
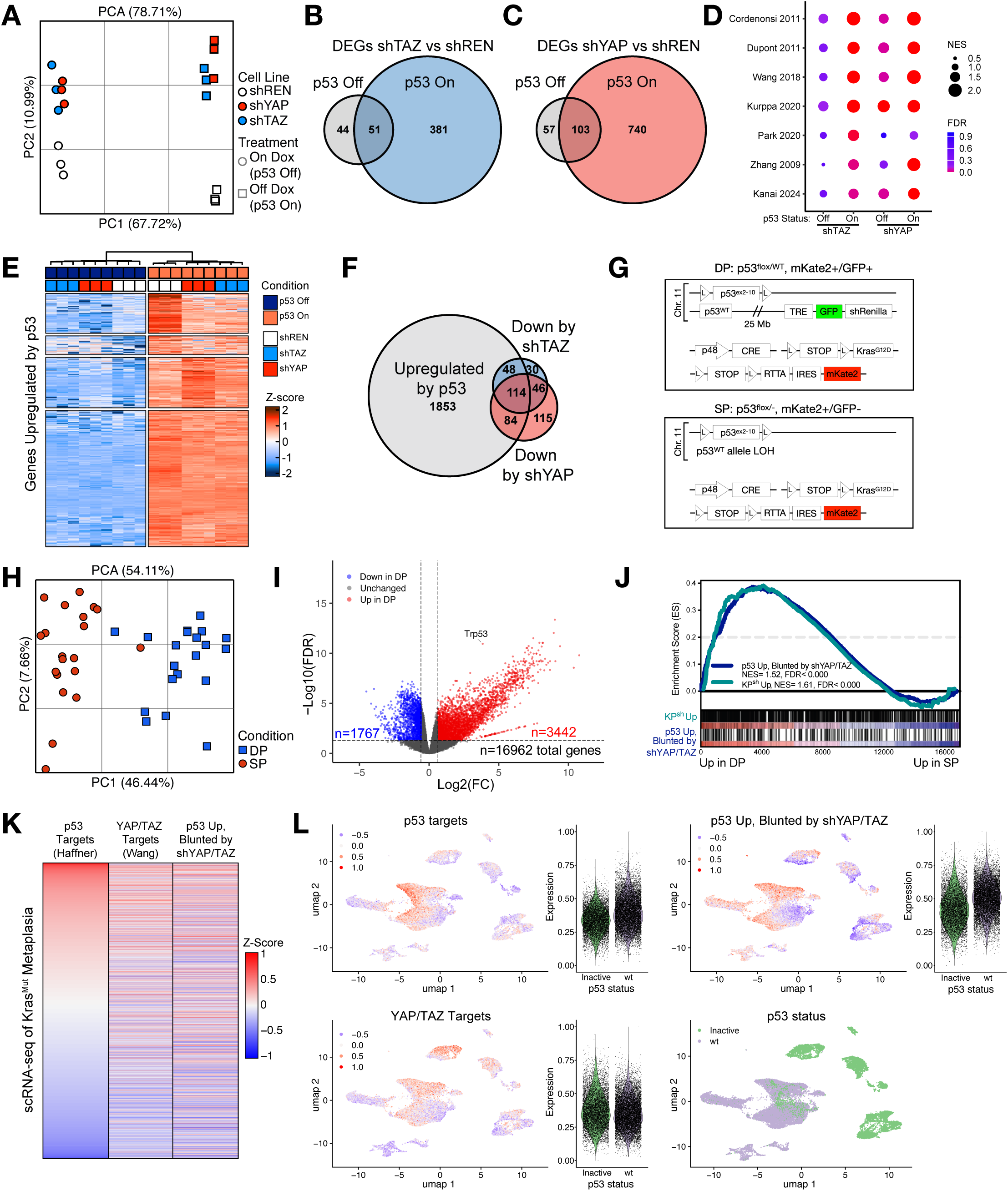
p53 determines transcriptional regulation by YAP/TAZ-TEAD. **(A-F)** RNA-seq from KP^sh^1 expressing shRNAs targeting Renilla luciferase (control, REN), Wwtr1 (TAZ), or Yap1 (YAP) on dox (p53 off) and 6 days off dox (p53 on). **(A)** PCA plot of all differentially expressed genes via ANOVA FC>1.5, FDR<0.05. **(B,C)** Venn diagrams of the number of genes with differential expression in **(B)** shTAZ vs shREN or **(C)** shYAP vs shREN depending on p53 status (DESeq2 FC>1.5, FDR<0.05). **(D)** Gene set enrichment analysis of published YAP/TAZ-TEAD gene signatures. **(E)** K-means clustered heatmap of all genes upregulated by p53 in shREN control cells (DESeq2 FC>1.5, FDR<0.05). **(F)** Venn diagram of genes in (E) overlapping those with decreased expression in shTAZ or shYAP cells with p53 on (DESeq2 FC<-1.5, FDR<0.05). **(G)** Allelic schematic of KPC^LOH^ model showing the pre-malignant p53 wild type GFP and mKate2 double positive (DP) and malignant p53 loss of heterozygosity mKate2 (SP) state. **(H-J)** Bulk RNA-seq of KPC^LOH^ mice from sorted DP and SP cell states. **(H)** PCA plot of DP and SP transcriptional states. **(J)** GSEA of genes upregulated in KP^sh^1 cells (KP^sh^ Up) and the KP^sh^ Up genes that are downregulated by depleting YAP or TAZ (p53 Up, Blunted by shYAP/TAZ). **(K-L)** scRNA-seq of Kras mutant pancreatic metaplasia from Burdziak 2023. **(K)** Heatmap correlation plot for p53 target genes (Haffner, 32070277), p53 Up, shYAP/TAZ blunted genes, and YAP/TAZ targets (Wang 2018) in neoplastic p53 wt pre-malignant pancreas cells. **(L)** UMAPs projecting the p53 cell state (K1-K4 for p53 wt and K5-K6 for p53 mut) and gene signature scores from (K) with adjacent violin plots of gene signature scores between cohorts stratified by p53 status.

Remarkably, RNA-seq analysis further revealed that YAP and TAZ function coordinates a substantial portion of p53 dependent gene expression. A majority of YAP and TAZ dependent genes were upregulated by restoring p53, with 68.1% (162/238) of genes downregulated by TAZ knockdown and 55.2% (198/359) of genes downregulated by Yap knockdown induced by p53 restoration (Fig. 2E,F). On the other hand, a smaller fraction of genes derepressed by Yap (∼30%, 60/194) or Taz (∼32%, 153/484) knockdown were downregulated by p53 restoration (Suppl. Fig. 4G,H). Yap or Taz knockdown blunted the expression of ∼13% of genes upregulated by p53 (246/1853, Fig. 2F) and increased expression of ∼10% of genes downregulated (169/1693, Suppl. Fig. 4H) revealing a previously uncharacterized ability of context dependent YAP/TAZ gene regulatory networks to shape global gene expression programs triggered by p53.

YAP and TAZ function are implicated in both the development of p53 wildtype, pre-malignant lineages as well as in maintenance of malignant progression following p53 inactivation during PDAC development^16,17,22^. Therefore, we asked if YAP/TAZ transcriptional targets licensed by p53 in KP^sh^ cells reflect stage-dependent expression patterns defined by p53 status in pancreatic cancer development. To determine global gene expression patterns linked with p53 in the PDAC pre-malignant to malignant transition, we took advantage of the KPC^LOH^ model. This model permits isolation of both Kras-mutant, pre-malignant cells where p53 remains intact and Kras-mutant malignant cells that have progressed following sporadic p53 loss of heterozygosity (LOH), via deletion of the remaining wildtype p53 allele and a cis-linked GFP expression cassette (Fig. 2G)^5^. We sorted cells based on p53 status (i.e., “double positive (DP)”: Kras^G12D^; p53^Δ/WT^; mKate+, GFP+ versus “single positive (SP)”: Kras^G12D^; p53^Δ/LOH^ (null); mKate+) from KPC^LOH^ mice following frank PDAC development. RNA-seq revealed extensive differences in gene expression in these two populations (Fig. 2H,I). Consistent with a role for p53 in licensing YAP/TAZ target gene expression, GSEA revealed enrichment of the genes upregulated by p53 restoration in KP^sh^ cells (KP^sh^ Up) as well as p53-triggered genes dependent on YAP or TAZ activity (“p53 Up, blunted by shYAP/TAZ”) in p53-intact, pre-malignant DP versus p53-inactivated SP PDAC cells (Fig. 2Jj). In addition, analysis of previously published single cell RNA-seq^13^ of Kras mutant, p53 wildtype pre-malignant cells (wt) versus p53 inactivated mouse PDAC (mut) revealed a strong correlation between p53 activity and p53 dependent YAP/TAZ targets in p53 wildtype, pre-malignant cells (Fig. 2K,L). Consistent with selective YAP/TAZ-TEAD regulation dictated by p53 activity, these p53-dependent YAP/TAZ targets displayed higher expression in p53-intact, pre-malignant versus p53-inactivated cancer transcriptional states, even compared with cancer cells where a high confidence YAP/TAZ target set derived from pan-cancer analysis was similarly expressed (Fig. 2L). Thus, p53 determines YAP/TAZ-TEAD target gene regulation that is predictive of stage-dependent expression patterns observed across the p53-dependent pre-malignant to malignant transition.

### Remodeled global occupancy in response to p53 activity primes distinct YAP/TAZ-TEAD transcriptional regulatory networks

Next, we asked if p53-dependent YAP/TAZ target regulation is determined by the redistribution of YAP/TAZ-TEAD binding in response to p53-triggered chromatin remodeling (Fig. 1). Annotating TEAD1, TEAD4, YAP, and TAZ peaks to genes upregulated by p53 that were blunted by Yap or Taz knockdown revealed increased local binding of all complex components, with significantly increased binding of at least one TEAD1, TEAD4, YAP, and TAZ peak at 52%, 42%, 35%, and 29% of genes, respectively (Fig. 3A). Increased YAP/TAZ-TEAD binding triggered by p53 corresponded with increased chromatin accessibility and elevated H3K27Ac at both local and remote putative enhancer regions of genes selectively regulated by YAP/TAZ, in a p53-dependent fashion, as exemplified by genes including Igfbp2 (within 10kb of the TSS, Fig. 3B) and Col11a1 (>50kb from the TSS, Suppl. Fig. 5A). We confirmed that upregulation of both Igfbp2 and Col11a1 following p53 restoration was YAP/TAZ-TEAD-dependent by showing that their upregulation was blocked by shRNAs targeting Yap or Taz or by the TEAD inhibitors VT104^32^ and IAG933^33^ (Fig. 3C,D and Suppl. Fig. 5B).

**Figure 3.**
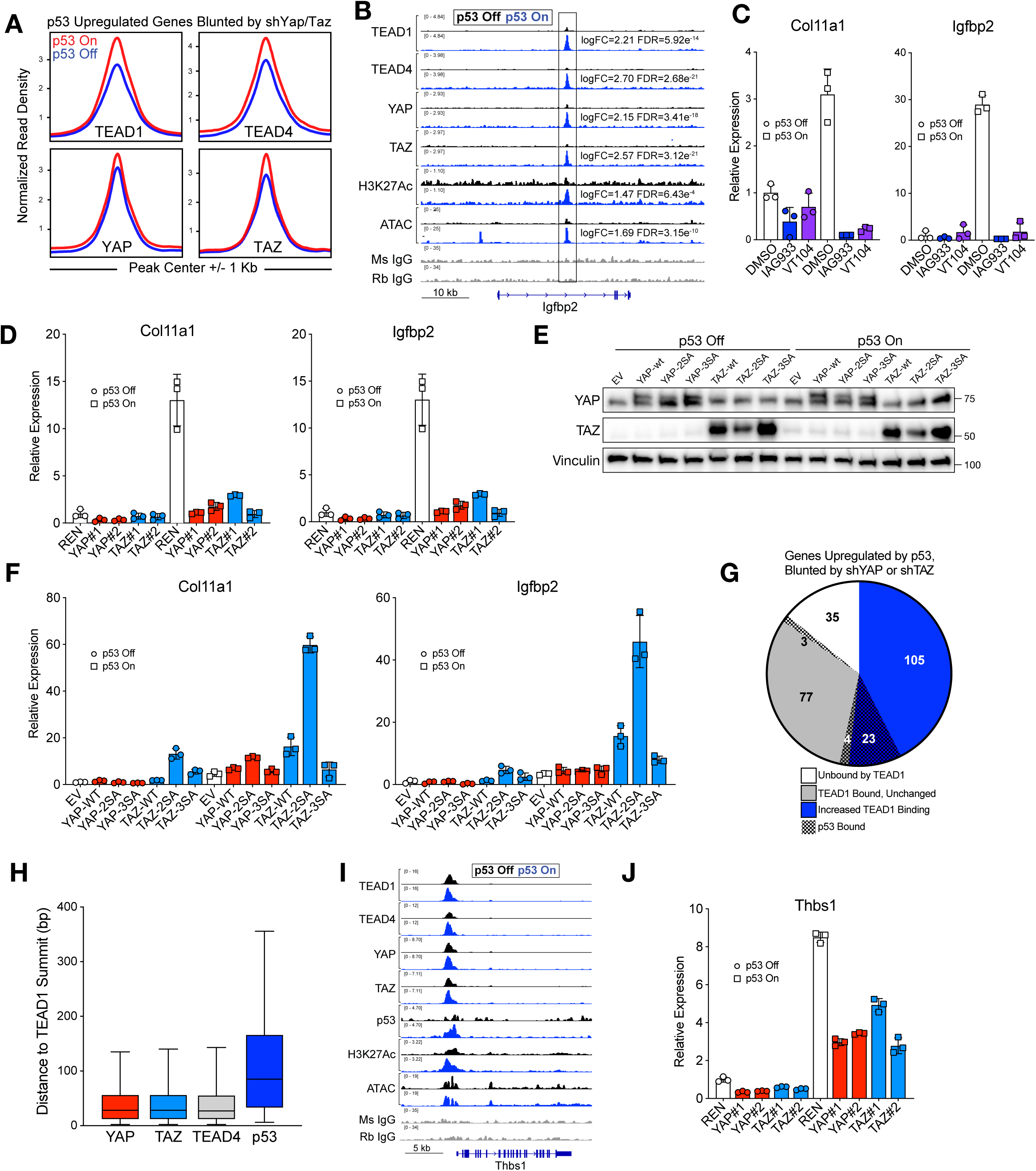
Redistribution of YAP/TAZ-TEAD primes p53-dependent transcriptional regulation. **(A)** Profile plots of normalized CUT&RUN peak density on dox (blue) or off dox (red) at TEAD1, TEAD4, YAP, and TAZ peaks at genes upregulated by p53 in shREN control cells and downregulated by shYAP or shTAZ in p53 on cells (FC>1.5, FDR<0.05). Peaks assigned to genes overlapped the gene body or were upstream of the TSS less than 5 Kb. **(B)** Normalized CUT&RUN and ATAC-seq signal in KP^sh^1 at the Igfbp2 locus. **(C-D)** RT-qPCRs KP^sh^1 **(C)** treated with 62 nM IAG933 or 500 nM VT104 or **(D)** expressing shREN, shYAP, or shTAZ upon 6 days of dox withdrawal normalized to Gusb. **(E-F)** KP^sh^1 were transduced with pBABE-Puro alone (EV) or various YAP/TAZ constructs. **(E)** Western blot of YAP/TAZ construct overexpression and **(F)** RT-qPCRs on dox (p53 off) or upon 6 days of dox withdrawal (p53 on) in KP^sh^1 normalized to Gusb. **(G)** Pie chart demonstrating variant TEAD1 binding or p53 binding at genes in (A). **(H)** Distances between peak summits overlapping a TEAD1 peak within 250 bp with boxes marking quartiles and whiskers denoting 95% of peaks. **(I)** Normalized CUT&RUN and ATAC-seq signal in KP^sh^1 at the Thbs1 locus. **(J)** RT-qPCR of Thbs1 in KP^sh^1 expressing shREN, shYAP, or shTAZ normalized to Gusb.

Taken together, these data suggest a model where p53 shapes chromatin landscapes and remodels TEAD binding, resulting in context-dependent transcriptional regulation of YAP/TAZ-TEAD complex targets in a p53 dependent fashion. To directly test this hypothesis, we asked if expression of p53-dependent YAP/TAZ-TEAD target genes would be disproportionately potentiated by increasing YAP or TAZ levels in KP^sh^ cells following p53 restoration. Thus, we constitutively overexpressed wildtype human YAP (YAP-wt) and TAZ (TAZ-wt) or phosphodeficient YAP S127A, S397A (YAP-2SA) and TAZ S89A, S311A (TAZ-2SA) which lack serine residues responsible for inhibitory phosphorylation and destabilization (Fig. 3E, Suppl. Fig. 5C)^23,34^. Consistent with p53-dependent TEAD binding acting to prime YAP/TAZ-dependent target gene expression, TAZ-wt and hyperactive TAZ-2SA potently induced expression of Igfbp2 and Col11a1 in KP^sh^ cells following p53 restoration while weakly inducing expression in p53-silenced cells (Fig. 3F, Suppl. Fig. 5D). Both genes were variably induced by YAP-wt and YAP-2SA, albeit to a lesser extent, likely due to lower levels of overexpression. YAP and TAZ mutants with an additional mutation preventing TEAD binding (YAP S94A (“YAP-3SA”) and TAZ S51A (“TAZ-3SA”)) failed to induce Col11a1 and Igfbp2 expression, demonstrating TEAD dependency of transcriptional regulation licensed by p53 activity (Fig. 3F, Suppl. Fig. 5D).

p53 acts as a sequence-specific transcription factor that results in transactivation of direct targets by binding response elements in promoter and early intronic regions^35^. Furthermore, p53 has been shown to cooperate with transcriptional effectors of other signaling pathways to drive specific targets via binding to distinct cis-binding elements at co-regulated target genes^36^. Thus, we integrated CUT&RUN profiling of p53 with YAP/TAZ-TEAD occupancy analysis to ask if direct, cooperative binding by YAP/TAZ-TEAD and p53 contribute to p53-dependent YAP/TAZ target gene expression. p53 CUT&RUN was robust, with strong correlation between replicates (Suppl. Fig. 5E), clear binding at well-characterized p53 target genes including Cdkn1a/p21 (Suppl. Fig. 5F), and enrichment of p53 binding motifs in identified peaks (Suppl. Fig. 5G).

Annotating TEAD1 peaks to genes upregulated by p53 in a YAP/TAZ dependent fashion revealed at least one TEAD1 peak detected at 84.6% (209/247) of genes, with 51.8% (128/247) of the genes displaying increased TEAD1 binding magnitude after restoring p53 (fold change>1.5, FDR<0.05). However, only 12.1% (30/247) displayed a p53 peak, of which 9.3% (23/247) exhibited both binding by p53 and increased magnitude of TEAD binding (Fig. 3G). p53 and YAP/TAZ-TEAD binding at these genes likely reflects engagement at distinct cis-regulatory elements rather than direct interactions at the chromatin, as the distance between summits of overlapping TEAD1-p53 peaks was more than 3 times that of overlapping TEAD1, YAP/TAZ, or TEAD4 peaks (85 vs 28, 28, and 27 base pairs respectively) (Fig. 3H). For example, offset binding of both p53 and YAP/TAZ/TEAD1/TEAD4 was detected at the promoter of the p53- and YAP/TAZ-dependent gene Thbs1, which was previously shown to be a target of both p53 and YAP/TAZ-TEAD at distinct sites^37,38^(Fig. 5I,J). Thus, while p53 and YAP/TAZ-TEAD directly coordinate some genes, p53 activity largely licenses YAP/TAZ-TEAD target selectivity through indirect effects on chromatin remodeling that primes targets for regulation by YAP/TAZ-TEAD complexes.

### p53 promotes TEAD target gene activation through actin-dependent TAZ accumulation

RNA-seq analysis of KP^sh^ cells expressing shRNAs targeting Yap or Taz revealed that p53 activity determines YAP/TAZ-TEAD target choice, but also the magnitude of their expression. (Fig. 4A, Suppl. Fig.4E). These data indicate that p53 activity not only remodels the repertoire of YAP/TAZ-TEAD targets but also potentiates the magnitude of TEAD dependent transactivation. Consistent with increased TEAD transcriptional activity in response to p53 restoration, GSEA revealed enrichment of both Hallmark p53 activation signatures as well as a pan-cancer YAP/TAZ-TEAD target gene signature^39^ in response to sustained p53 restoration (Fig. 4A, Suppl. Fig. 6A).

**Figure 4:**
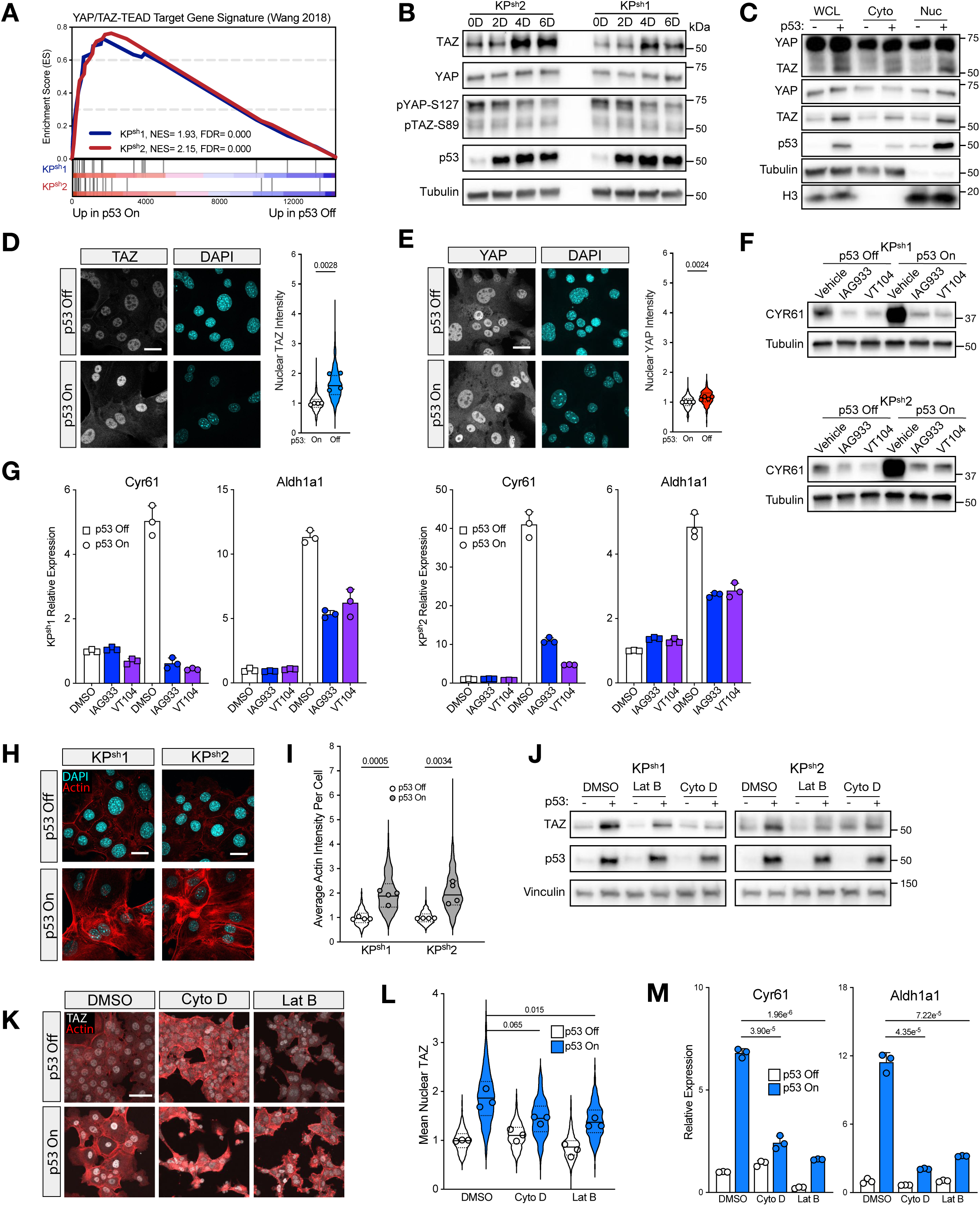
p53 promotes TEAD activation through actin-induced TAZ accumulation. **(A)** GSEA analysis of YAP/TAZ-TEAD target gene signature (Wang 2018) from RNA-seq of KP^sh^1 and KP^sh^2 cells after 6 days of dox withdrawal (p53 on). **(B)** Western blot was performed on indicated days (0, 2, 4, and 6) following doxycycline withdrawal. **(C)** Western blots on whole cell lysate (WCL), Cytoplasm (Cyto), and Nuclear (Nuc) KP^sh^1 lysate cell fractions on dox (p53 -) and 6 days off dox (p53 +). **(D-E)** Immunofluorescence microscopy images of KP^sh^1 subjected to 6 days of dox withdrawal (p53 on) and quantification of nuclear **(D)** TAZ and **(E)** YAP staining intensity. Dots show median signal across separate wells of independent experiments and violin shows distribution combined across all experiments. p-value obtained using student’s t-test comparing medians across experiments. **(F)** Western blots and **(G)** RT-qPCR of KP^sh^1 and KP^sh^2 on dox (p53 off) or 6 days off dox (p53 on) treated with 48 hours of 1:1000 DMSO, 62 nM IAG933, or 500 nM VT104. **(H)** Fluorescence microscopy images of KP^sh^1 and KP^sh^2 upon 6-day dox withdrawal (p53 on) stained with Phalloidin (Actin) and DAPI and **(I)** associated quantification from four independent experiments. Dots show median signal across separate wells of independent experiments and violin shows distribution combined across all experiments. p-value obtained using student’s t-test on median cellular actin intensity across experiments. **(J-M)** KP^sh^1 treated with 12 hours of actin polymerization inhibitors Latrunculin B (Lat B, 5 µM) or Cytochalasin D (Cyto D, 1 µM) compared to vehicle (1:1000 DMSO) on dox (p53 off) or 6 days of dox withdrawal (p53 on). **(J)** Representative western blots, **(K)** IF microscopy images, and **(L)** quantification of IF microscopy images with dots showing median signal of three independent experiments and violin showing distribution combined across all experiments in KP^sh^1. p-value obtained using student’s t-test of median nuclear signal from three experiments. **(M)** RT-qPCR normalized to Gusb, p-value from student’s t-test. Scale bars = 25 µm in (C,D,H) and 50 µm in (K).

While p53 restoration did not increase TEAD protein levels (Suppl. Fig. 6B), it triggered progressively increased levels of TAZ (Fig. 4B), providing a mechanism for increased TEAD target gene expression. Phosphorylation of YAP/TAZ promotes cytoplasmic sequestration and initiates ubiquitin-mediated degradation^40^. Accordingly, p53 restoration resulted in decreased levels of phosphorylated YAP/TAZ, corresponding with notable increases in total cellular and nuclear TAZ levels and with a modest increase in nuclear YAP (Fig. 4B-E; Suppl. Fig. 6C-E). Neither Yap nor Taz mRNA expression was elevated by p53 restoration, suggesting their levels are regulated post-translationally (Suppl. Fig. 6F). Like shRNA-mediated silencing of Yap and Taz, disrupting YAP/TAZ-TEAD interactions by small molecule inhibition prevented upregulation of TEAD target genes Cyr61 and Aldh1a1, implicating TAZ coactivation of TEAD in elevated TEAD target gene expression following p53 restoration (Fig. 4F,G).

YAP/TAZ levels frequently reflect activity of the Hippo signaling cascade, which promotes YAP/TAZ phosphorylation and degradation through MST1/2 kinase-mediated phosphorylation and activation of the LATS1/2 kinases, enhanced by the MOB1 scaffolding protein 40. However, despite decreased YAP/TAZ phosphorylation, p53 restoration decreased neither phosphorylation nor expression of LATS1, LATS2, or MOB1, suggesting that a Hippo-independent mechanism drives TAZ stabilization and nuclear accumulation (Suppl. Fig. 6G). YAP/TAZ levels can be regulated independently of Hippo signaling in response to changes in F-actin cytoskeleton tension, organization, and abundance^41^. Indeed, visualization of the actin cytoskeleton in response to p53 restoration revealed a striking reorganization of F-actin, including stress fiber polymerization and elevated actin abundance that is consistent with morphological expansion and increased cell size^42,43^ (Fig. 4 H,I). To test if actin polymerization promotes TAZ accumulation in response to p53 restoration, we treated KP^sh^ cells with inhibitors of actin polymerization, Latrunculin B (Lat B) and Cytochalasin D (Cyto D). Inhibiting actin polymerization with both agents blunted accumulation of TAZ protein and reduced its nuclear accumulation in response to sustained p53 restoration (Fig. 4J-L and Suppl. Fig. 6H,I). Furthermore, inhibition of actin polymerization was sufficient to blunt the upregulation of the TEAD targets Cyr61 and Aldh1a1 upon p53 restoration (Fig. 4M and Suppl. Fig. 6J). Taken together, these results demonstrate that p53 activity can determine the magnitude of TEAD target gene expression by increasing TAZ levels in response to altered cellular actin dynamics.

### Actin reorganization in response to cell cycle arrest and cellular senescence connects p53 activity to TAZ accumulation

We next set out to determine the mechanism connecting p53 activity to actin-mediated TAZ accumulation. While p53 can directly regulate genes associated with actin coordination^42^, accumulation of actin stress fibers is also a common feature of cell cycle arrest and cellular senescence^42,43^, triggered by sustained p53 restoration in KP^sh^ cells (Suppl. Fig. 1C,D). Therefore, to ask if p53 activation in the absence of cell cycle arrest is sufficient to remodel actin and trigger TAZ accumulation, we knocked down Cdkn1a/p21, a key p53 direct target that enforces cell cycle arrest and senescence^45^. Stable p21 knockdown maintained Ki67+ proliferation and significantly blunted induction of senescence-associated β-galactosidase (SA-β-Gal) despite p53 accumulation in response to dox withdrawal from KP^sh^ cells (Fig. 5A,B; Suppl. Fig. 7A,B). Furthermore, p21 knockdown prevented the increases in cellular TAZ levels and nuclear accumulation observed in response to p53 restoration (Fig. 5B-D; Suppl. Fig. 7C-E), and corresponding with reduced actin accumulation (Fig. 5E; Suppl. Fig. 7F). Therefore, we nominate actin reorganization as a stress response associated with cell cycle arrest and cellular senescence as a key cell fate change which connects p53 activity to increased TAZ levels and increased TEAD target gene expression.

**Figure 5.**
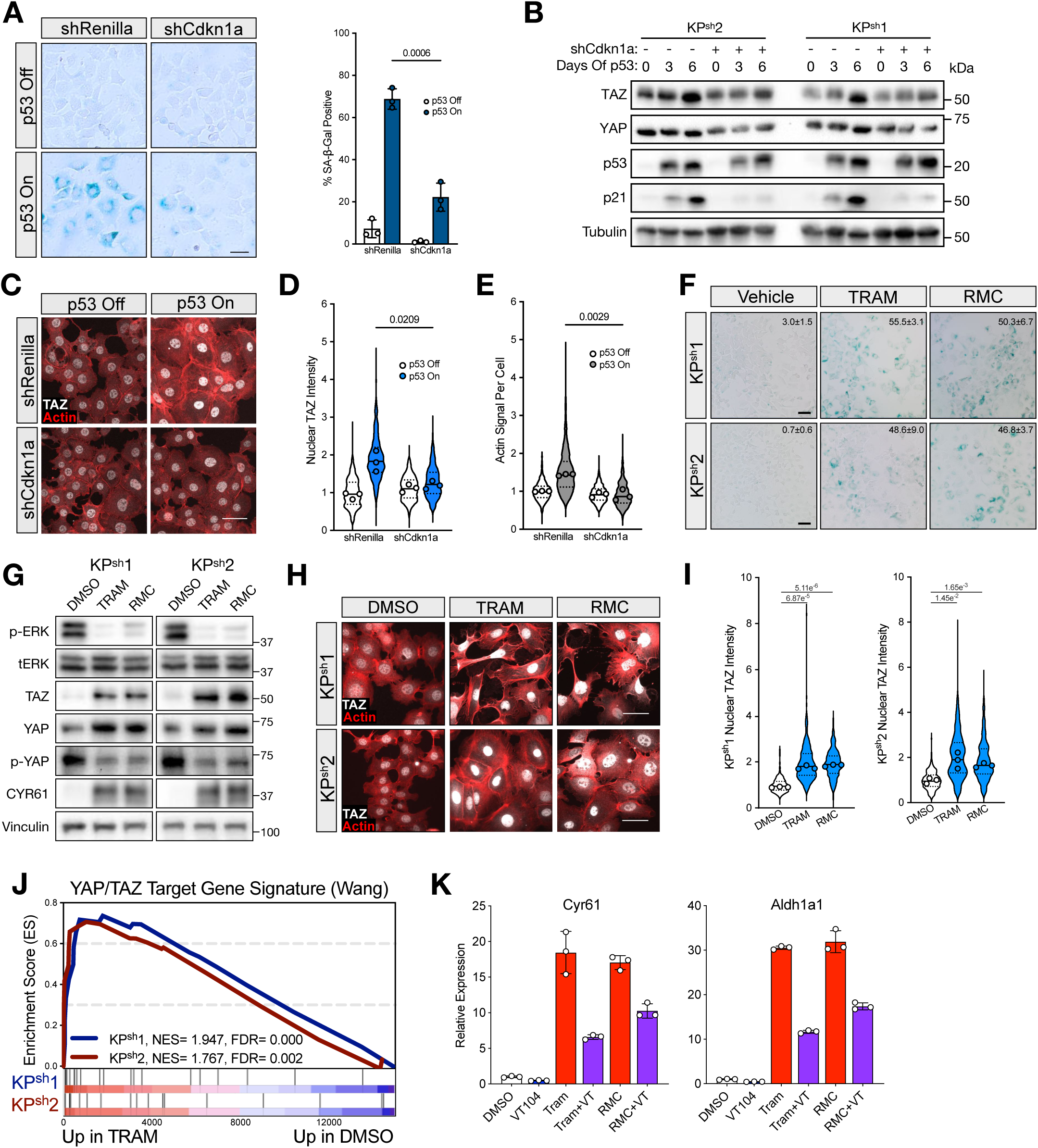
Senescence-induced actin reorganization induces TAZ accumulation. **(A-E)** KP^sh^1 and KP^sh^2 cells expressing shRNA targeting Renilla luciferase (control) or Cdkn1a. **(A)** Representative SA-β-Galactosidase images of KP^sh^1 (left) and quantification of three independent wells (right). p-value from student’s t-test. **(B)** Western blot upon dox withdrawal at indicated time points. **(C)** IF microscopy of TAZ and Phalloidin staining upon restoring p53 with associated quantification of **(D)** nuclear TAZ and **(E)** total Actin per cell in KP^sh^1. Dots show median signal across separate wells of independent experiments and violin shows combined distribution from all experiments. p-value obtained using student’s t-test of median signal from triplicate experiments. **(F-I)** KP^sh^ were treated with 72 hours of 50 nM Trametinib, 50 nM RMC, or DMSO vehicle matched control**. (F)** Representative SA-β-Gal images with quantification of three independent wells with percent SA-β-Gal positive cells and standard deviation in upper right corner of images. **(G)** Western blots and **(H)** representative IF images of TAZ and Phalloidin (Actin) with **(I)** associated quantification of KP^sh^1 and KP^sh^2 nuclear TAZ. Dots show median signal across three separate wells and violin shows combined distribution of all wells. p-value obtained using student’s t-test of median signal from triplicate wells. **(J)** GSEA of RNA-seq from KP^sh^ treated with 72 hours of 25 nM Trametinib or matched DMSO vehicle. **(K)** RT-qPCRs from KP^sh^1 treated with 72 hours of 50 nM Trametinib, 50 nM RMC, or DMSO then an additional 48 hours with or without the addition of 500 nM VT104. Scale bars = 50 µm in (A, C, H) and 100 µm in (F).

We next asked if cell cycle arrest and senescence in the absence of p53 activity was sufficient to promote TAZ activation. Inhibition of Ras/MAPK signaling drives cell cycle arrest and senescence in p53-silenced KP^sh^ cells^15^. Therefore, we employed the MEK inhibitor trametinib (TRAM) and the pan-RAS inhibitor RMC-7977 (RMC) to ask if pharmacologically induced arrest and senescence could increase actin and TAZ levels and elevate TEAD target gene expression in the absence of p53 accumulation. TRAM and RMC effectively decreased ERK phosphorylation, an essential effector of KRAS-driven MAPK signaling^46^, reduced Ki67+ proliferation, and triggered SA-β-Gal positivity in p53-silenced KP^sh^ cells (Fig. 5F,G; Suppl. Data Fig. 7G). Arrest and induction of senescence markers corresponded with robust TAZ accumulation, a modest increase in total YAP accompanied by decreased phosphorylation, and increased levels of the YAP/TAZ-TEAD target CYR61 (Fig. 5G). Activation of TAZ corresponded with elevated levels of actin as observed in response to p53 restoration (Fig. 5H,I). As in the response to sustained p53 activity, RNA-seq analysis of p53-silenced KP^sh^ cells treated with TRAM revealed enrichment of YAP/TAZ-TEAD target genes (Fig. 5J) coincident with depletion of proliferation-associated gene sets indicative of arrest and senescence (e.g., Hallmark E2F targets, Myc targets (Suppl. Fig. 7H). Accordingly, induction of senescence via TRAM or RMC followed by co-treatment with the TEAD inhibitor VT104 blunted upregulation of YAP/TAZ-TEAD target genes Cyr61 and Aldh1a1 (Fig. 5K; Suppl. Data Fig. 7I). Taken together, these data demonstrate that actin remodeling associated with cell cycle arrest and senescence triggers TAZ accumulation to drive TEAD-mediated gene transactivation, representing a mechanism by which cell fates induced by p53 can connect with transcriptional regulation by YAP/TAZ-TEAD.

### YAP/TAZ-TEAD regulation licensed by p53 activity coordinates senescence-associated secretory phenotypes

We next investigated the expression programs controlled by p53 licensed YAP/TAZ-TEAD transcriptional regulation. Ontology overrepresentation analysis of genes upregulated by p53 in a YAP/TAZ dependent fashion revealed enrichment of gene sets associated with remodeling of cellular microenvironments and epithelial to stromal communication (e.g., angiogenesis and extracellular matrix organization, Fig. 6A). A hallmark feature of p53-induced senescence is secretion of an array of proteins including growth factors, cytokines, proteases, and extracellular matrix components broadly classified as the Senescence-Associated Secretory Phenotype (SASP)^47^, prompting us to ask if p53-licensed YAP/TAZ-TEAD activity regulates SASP expression. GSEA validated that p53 restoration in KP^sh^ cells induced robust upregulation of SASP genes (“SASP ATLAS” gene set, Fig. 6B) and, remarkably, that SASP expression displayed p53-dependent, selective regulation by YAP/TAZ, as the SASP signature was significantly depleted only upon Yap or Taz knockdown in KP^sh^ cells following p53 restoration (Fig. 6B). This SASP signature reflects coordinated expression of cytokines, growth factors, insoluble collagens, and the Loxl family of collagen crosslinking enzymes (Suppl. Fig. 8A).

**Figure 6:**
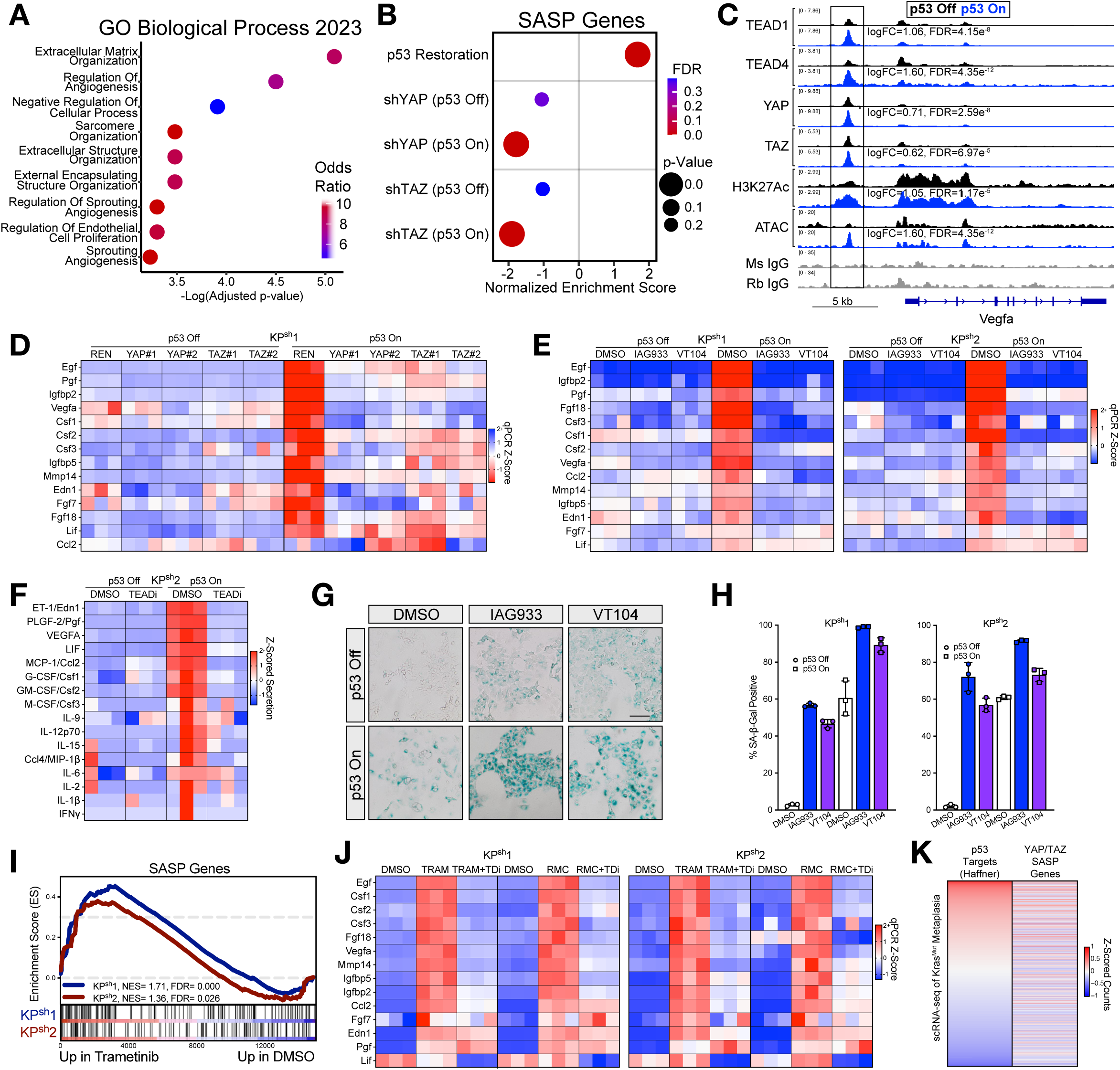
YAP/TAZ-TEAD transactivates SASP genes through enhancer binding during senescence. **(A)** Overrepresentation analysis (Enrichr, GO Biological Processes 2023) of genes upregulated by p53 in KP^sh^1 shREN cells and downregulated by shYAP or shTAZ in p53 on cells (FC>1.5, FDR<0.05). **(B)** Gene set enrichment analysis of SASP Atlas genes. **(C)** Normalized CUT&RUN and ATAC-seq tracks at the Vegfa locus. **(D,E)** RT-qPCR for genes of secreted proteins related to the SASP in **(D)** KP^sh^1 with shREN, shYAP or shTAZ and **(E)** KP^sh^1 and KP^sh^2 treated with 48 hours of 1:1000 DMSO, 62 nM IAG933, or 500 nM VT104 on dox (p53 off) and 6 days off dox (p53 on) normalized to Gusb. **(F)** Z-scored concentration of cytokines secreted from KP^sh^2 treated with 1:1000 DMSO or 62 nM IAG933 between 24-72 hours on dox (p53 off) or 6 days off dox (p53 on). **(G)** Representative SA-β-Galactosidase images of KP^sh^1 treated with 48 hours of 1:1000 DMSO, 62 nM IAG933, or 500 nM VT104. **(H)** Quantification of percent SA-β-Galactosidase positive cells from (G). **(I)** GSEA of RNA-seq from KP^sh^ cells treated with 72 hours of 25 nM Trametinib. **(J)** RT-qPCRs from KP^sh^ treated with 72 hours of 50 nM Trametinib, 50 nM RMC, or DMSO followed by an additional co-treatment with 48 hours of 500 nM VT104. Separate Z-scores were calculated for TRAM and RMC treatment. **(K)** Heatmap correlation plot of scRNA-seq data from Burdziak 2023 for the p53 wt pre-malignant cell state showing how YAP/TAZ regulated SASP genes from Figure 7D correlate with p53 target gene enrichment (Haffner, 32070277).

p53-licensed regulation of secreted factors by YAP/TAZ-TEAD was dependent on the two levels of molecular interaction between p53 activity and YAP/TAZ-TEAD described above (i.e., priming of targets via increased TEAD binding and elevated Taz levels). For example, several genes encoding secreted proteins licensed by p53, including the pro-angiogenic factors Vegfa and Pgf, displayed increased binding of TEAD1, TEAD4, YAP, and TAZ at putative enhancers associated with p53-triggered increases in H3K27Ac abundance and chromatin accessibility (Fig. 6C; Suppl. Data Fig. 8B). Other upregulated SASP genes (e.g., Csf3/G-CSF) were bound by TEAD1, TEAD4, YAP, and TAZ independently of p53 activity but displayed increased expression coincident with elevated Taz (Suppl. Data Fig. 8C). Yap/Taz knockdown and pharmacological inhibition of TEAD selectively blocked expression of both classes of SASP factors in response to p53 restoration (Fig. 6D,E) while their secretion was selectively blocked by inhibiting TEAD with IAG933 (Fig. 6F, Suppl. Fig. 8B,C). Taken together, p53 activity expands YAP/TAZ-TEAD regulation of secreted factors, both by increased YAP/TAZ-TEAD complex binding and by potentiating TEAD transactivation largely through TAZ accumulation.

YAP/TAZ-TEAD activity has been shown to contextually drive or impair senescence^48^. Demonstrating that TEAD inhibition in KP^sh^ cells reduced SASP gene expression by blocking its transcriptional activity rather than by impairing senescence induction, treatment with VT104 or IAG933 increased SA-β-Gal staining in response to p53 restoration (Fig. 6G,H). Consistent with cell cycle arrest and senescence establishing YAP/TAZ-TEAD-dependent regulation of secretory programs, TRAM treatment resulted in SASP signature enrichment in KP^sh^ cells where p53 remained silenced (Fig. 6I), and in increased expression of secretedfactors largely overlapping with those upregulated in a YAP/TAZ-TEAD dependent fashion in response to p53 restoration (Suppl. Data Fig. 8D). Small molecule inhibition of TEAD transcription following TRAM or RMC treatment was sufficient to blunt upregulation of most of these secreted factors, thus confirming TEAD dependency in this context (Fig. 6J).

Like the broader set of YAP/TAZ-TEAD targets licensed by p53 activity in KP^sh^ cells (“p53 Up, blunted by shYAP/TAZ”, Fig. 2F,K,L), analysis of single-cell expression data from p53-intact pre-malignant cells and p53-inactivated PDAC cells reveals a direct correlation between p53 targets and p53-licensed YAP/TAZ-TEAD-regulated secretory genes (Fig. 6K). This includes secreted factors that play prominent roles in establishing immune suppressive microenvironments that promote pre-malignant specification (e.g., GM-CSF/Csf2)^49,50^ as well as determinants of tumor-associated vasculature (e.g., Vegfa)^51^ (Fig. 7D-F). Thus, licensing of YAP/TAZ-TEAD transcriptional regulation by p53, in part through cellular adaptations to cell cycle arrest and senescence, determines distinct secretory programs that are implicated in epithelial to stromal communication in pancreatic cancer and may play a role in the establishment of stage-dependent microenvironments during PDAC progression.

## DISCUSSION

Stepwise carcinogenesis is characterized by emergent transcriptional plasticity and novel epigenetic states that are accessed as transforming cells acquire collaborating cancer-driving mutations^52^. Epigenetic remodeling in cancer results in “hijacking” of transcription factors that otherwise regulate normal development and homeostasis to drive novel expression programs that establish neoplastic cell fates^53,54^. As the most frequently mutated gene in cancer, inactivation of the tumor suppressor p53 represents the most common genetic route by which malignant biology emerges, frequently in the setting of the pre-malignant to malignant switch, but the mechanisms that connect p53 activity to distinct transcriptional programs during cancer development remain poorly understood. Here, we nominate coordination by p53 of both the regulatory networks and the transcriptional activity of YAP/TAZ-TEAD complexes as a novel determinant of expression landscapes in pancreatic cancer progression, a tumor type where p53 mutation represents a dominant route to malignant progression of neoplastic cells initiated by oncogenic Kras. We leverage a genetically engineered mouse model of PDAC which permits restoration of endogenous p53 function in malignant cells, representing a unique platform to identify context dependent effects of p53 activity on chromatin regulatory landscapes in an isogenic setting. As previously demonstrated^15^, restoring p53 activity in PDAC cells triggers features of pre-malignant gene expression, and here we show that this reflects remodeled chromatin accessibility landscapes associated with TF recognition motifs that overlap with those gained and lost in the pre-malignant to malignant shift triggered by p53 inactivation^13,14^. Sustained p53 restoration results in profound effects on chromatin accessibility as well as on histone modifications that define the activity of regulatory regions including promoters and enhancers, which we link to context-dependent occupancy and transcriptional regulation of YAP/TAZ-TEAD triggered by p53 activity. Thus, our data provides novel insight into how p53 activity guides the emergence of transcriptional hierarchies during cancer development, where chromatin landscapes dependent on p53 activity influence the regulatory networks and transcriptional outputs of targets of key transcription factors, like YAP/TAZ-TEAD, in a stage-dependent fashion.

YAP/TAZ-TEAD complexes play a critical role in normal tissue development and homeostasis^55^ and represent a prototypical example of a transcriptional regulator that is aberrantly regulated during stepwise pancreatic tumorigenesis. While YAP and TAZ accumulate during pancreatic regeneration^16^, their levels remain elevated during the development of p53 wildtype, pre-malignant ductal metaplasia and pancreatic intraepithelial neoplasia (PanIN) triggered by mutant Kras^16,21^ and in Kras mutant, p53 inactivated malignancy^17^. However, YAP/TAZ-TEAD regulatory networks at each stage remain incompletely defined. We describe a model (Suppl. Data Fig. 8E) by which p53 activity can determine, at two levels of molecular control, the transcriptional outputs of YAP/TAZ-TEAD in cells transformed by mutant Kras. First, we demonstrate that p53 activity alters chromatin landscapes that expand the repertoire of YAP/TAZ-TEAD occupancy, connecting YAP/TAZ-TEAD to a distinct set of transcriptional targets when p53 is engaged. Consistent with YAP/TAZ-TEAD largely regulating gene expression at active enhancers^56^, YAP/TAZ-TEAD binding at targets licensed by p53 activity reflects increased enhancer chromatin accessibility and activity. Thus, our work nominates YAP/TAZ-TEAD as an interpreter of dynamic enhancer landscapes triggered by p53 activity, including higher order enhancer architecture, as TEAD motifs represent the most enriched motif in super-enhancers opened upon p53 restoration. Taken together, regulation of a broader set of YAP/TAZ-TEAD target genes in cells with active p53 adds novel detail to the consequence of p53 activity on global transcriptional regulatory landscapes ^57^. Critically, YAP/TAZ-TEAD targets licensed by p53 activity through chromatin remodeling identified in KP^sh^ cells display stage-dependent expression in genetically engineered models of PDAC where YAP/TAZ-TEAD signaling is heterogeneous across both p53-intact pre-malignant and p53-inactivated malignant populations. YAP and TAZ levels depend on varied signaling and physiological cues that shape PDAC progression^58^ and this work suggests that p53 acts to determine the diversity of TEAD targets regulated by these cues in a stage-dependent fashion.

Our work also identifies actin-dependent accumulation of TAZ in response to p53-induced arrest and cellular senescence as one of these putative physiological cues, representing is a second level of molecular control connecting p53 activity to transcriptional regulation by TEAD (Suppl. Data Fig. 8e). Stress fibers accompanying actin reorganization are a prominent feature of p53-triggered senescence^43,59^, and our model extends actin levels from a regulator of senescent morphology to a functional signaling node that determines TAZ levels and TEAD target gene expression licensed by p53 activity. Here, senescence established by sustained p53 activity, or through other triggers including inhibition of the KRAS/MAPK pathway, results in rapid cellular accumulation and nuclear translocation of TAZ and thus in selective TEAD target gene activation. Notably, while YAP and TAZ phosphorylation and degradation are frequently regulated by the HIPPO signaling pathway, we observe no change in the levels or activity of the upstream LATS1/2 kinases in response to p53 restoration, suggesting HIPPO-independent regulation of TAZ levels. This is consistent with the actin-mediated control of TAZ observed here, and in both Drosophila and mammalian cells where actin-mediated control of YAP and TAZ levels does not require LATS kinase function^60^.

Remarkably, both levels of p53 control of YAP/TAZ-TEAD activity converge on cellular adaptation to cell cycle arrest and senescence. Chromatin remodeling is a hallmark of both replicative and oncogene-induced senescence, entailing intricate remodeling of enhancer landscapes that are implicated in the coordination of key biological features of senescence, including the SASP^28,61^. Accordingly, remodeling of chromatin accessibility and enhancer landscapes that correspond with reshaping of YAP/TAZ-TEAD genomic occupancy in response to p53 activity occurs coincident with cell cycle arrest and markers of senescence in vitro. Furthermore, senescence is a hallmark of Kras mutant, p53 wildtype, pre-malignant cells^62,63^, which display selective expression of p53-licensed YAP/TAZ-TEAD targets. Arrest and senescence triggered by potent inhibition of the Kras/MAPK pathway in p53-inactivated cells also results in accumulation of TAZ and confers TEAD dependent control of secreted proteins that are also licensed by p53 activity. Thus, our work nominates YAP/TAZ-TEAD complexes as novel transcriptional regulators of senescence-associated transcriptional programs, including the SASP. Notably, because expression of SASP genes in models of replicative senescence are associated with exclusion, rather than binding, of TEAD at open chromatin flanking secreted genes^64^, our work suggests that TEAD mediated regulation of SASP genes may also be associated with senescence in oncogene expressing cells.

In summary, this work elucidates mechanisms by which p53 activity shapes transcriptional plasticity during tumorigenesis by diversifying the outputs of transcription factors involved in establishing aberrant neoplastic cell fate. We deeply characterize the ability of p53 activity to license target choice and expression of YAP/TAZ-TEAD complexes and highlight the role whereby cellular adaptation to p53-triggered changes in cell biology, e.g., cell cycle arrest and senescence, reverberates on transcriptional programs defined by p53 status. Furthermore, as TEAD inhibition is emerging as a potential therapeutic strategy in many tumor types including PDAC^23,65^, , our work suggests that the response to TEAD inhibitors may depend on p53 activity, or induction of cell fates associated with p53 activity, such as senescence, that can be induced by putative combination therapies, including newly emerging RAS inhibitors^23,66^.

## Supplementary Figure Legends

**Supplementary Figure 1:**
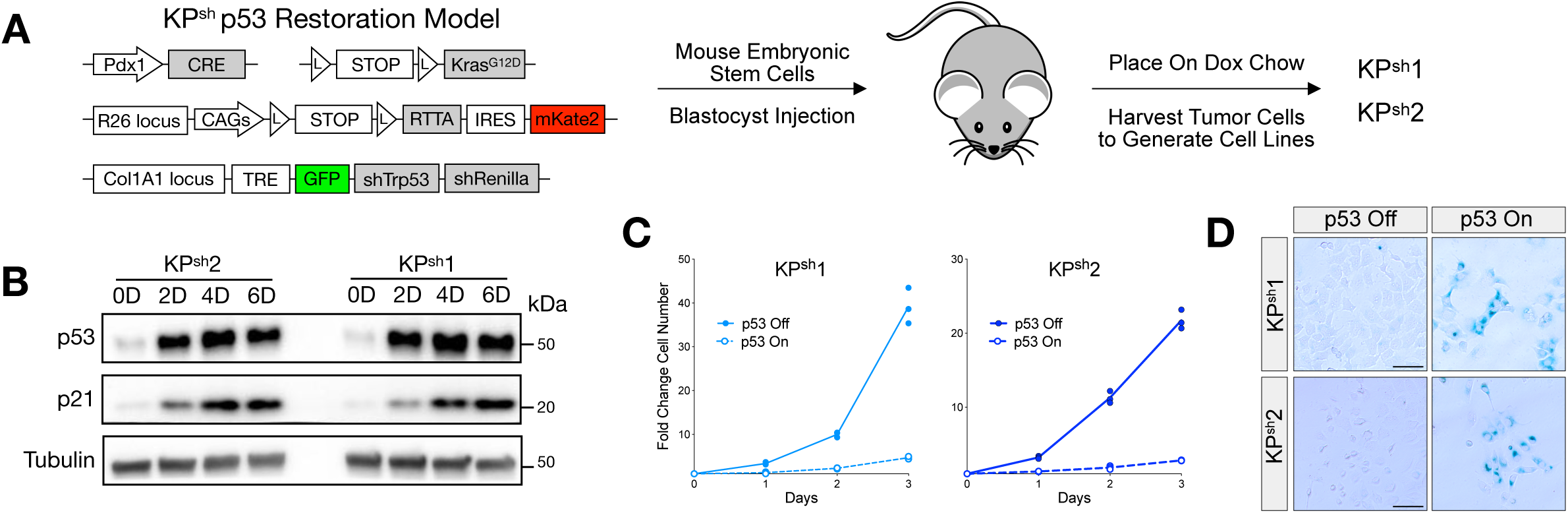
Restoration of wildtype p53 in a mouse model of pancreas cancer. **(A)** Schematic showing generation of the <u>K</u>ras^G12D^, doxycycline (dox) inducible <u>p</u>53 <u>sh</u>RNA (KP^sh^) pancreas cancer model. **(B)** Western blot of time-dependent p53 restoration upon doxycycline withdrawal in two independently derived KP^sh^ cell lines. **(C)** Growth curve of KP^sh^1 and KP^sh^2 on dox (p53 off) or measured daily beginning after 2 days of dox withdrawal (p53 on). **(D)** Representative SA-β-Galactosidase images of KP^sh^1 and KP^sh^2 on dox (p53 off) or after 6 days of doxycycline withdrawal (p53 on). Scale bar = 100 µm in (D).

**Supplementary Figure 2:**
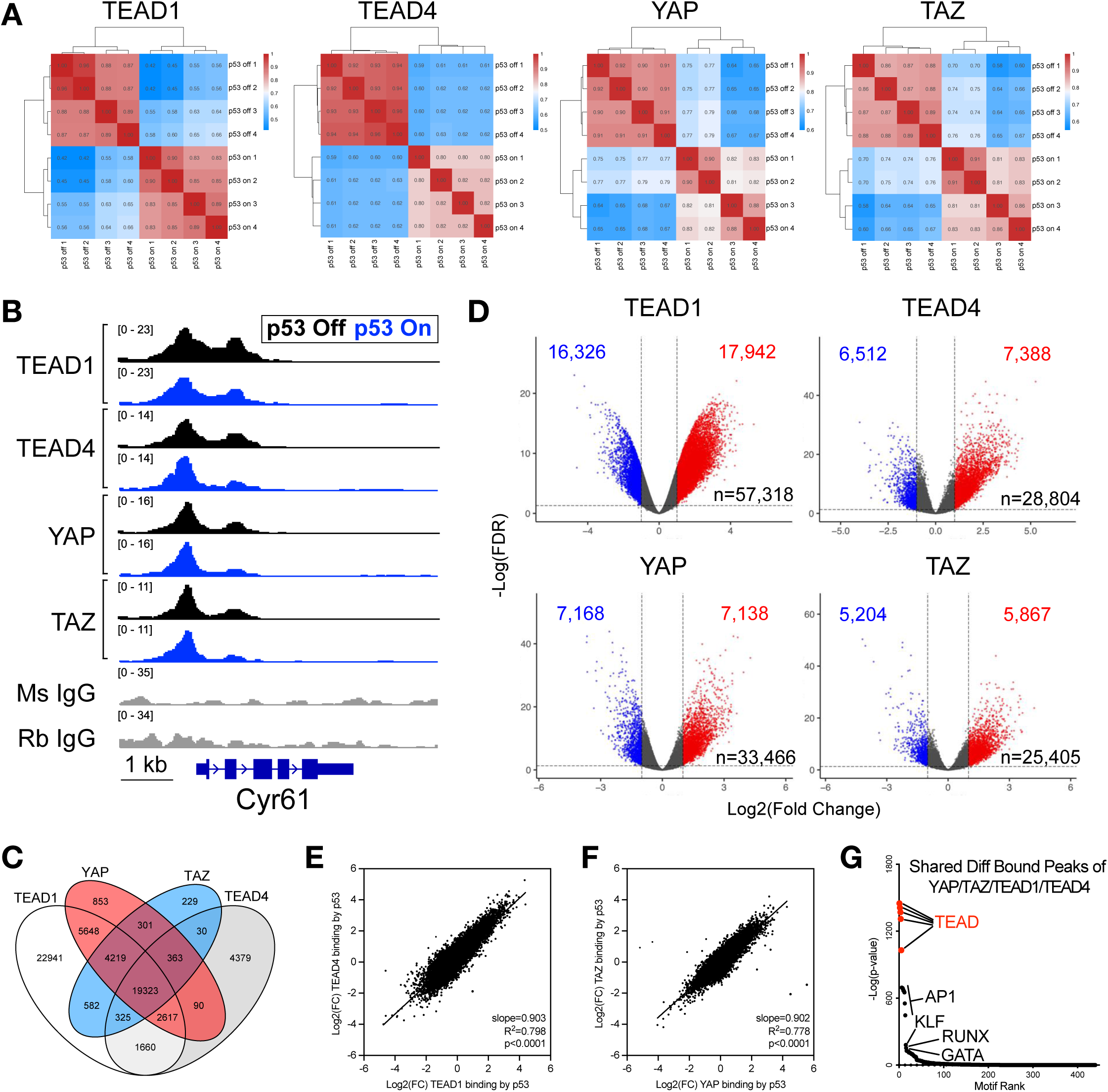
YAP/TAZ-TEAD1/4 exhibit overlapping genomic occupancy redistribution upon p53 restoration. **(A-G)** CUT&RUN was performed in KP^sh^1 cells on dox (p53 off) and 6 days following dox withdrawal (p53 on) for TEAD1, TEAD4, YAP, TAZ, and isotype matched control IgGs. **(A)** Heatmaps of binding affinity correlation across quadruplicates plates collected for CUT&RUN. **(B)** Normalized average tracks at the YAP/TAZ-TEAD target gene Cyr61. **(C)** Venn diagram of overlapping peaks called in TEAD1, TEAD4, YAP, and TAZ CUT&RUN. **(D)** Volcano plots of differential binding upon p53 restoration (EdgeR) with dots colored in red and blue denoting peaks with significantly increased and decreased binding (FC>1.5, FDR<0.05). **(E-F)** Correlation plot of differential binding magnitude upon restoring p53 in **(E)** TEAD1 vs TEAD4 and **(F)** YAP vs TAZ. **G)** Homer known motif analysis on all TEAD1, TEAD4, YAP, and TAZ peaks with differential binding detected in all four proteins from Figure 1e.

**Supplementary Figure 3:**
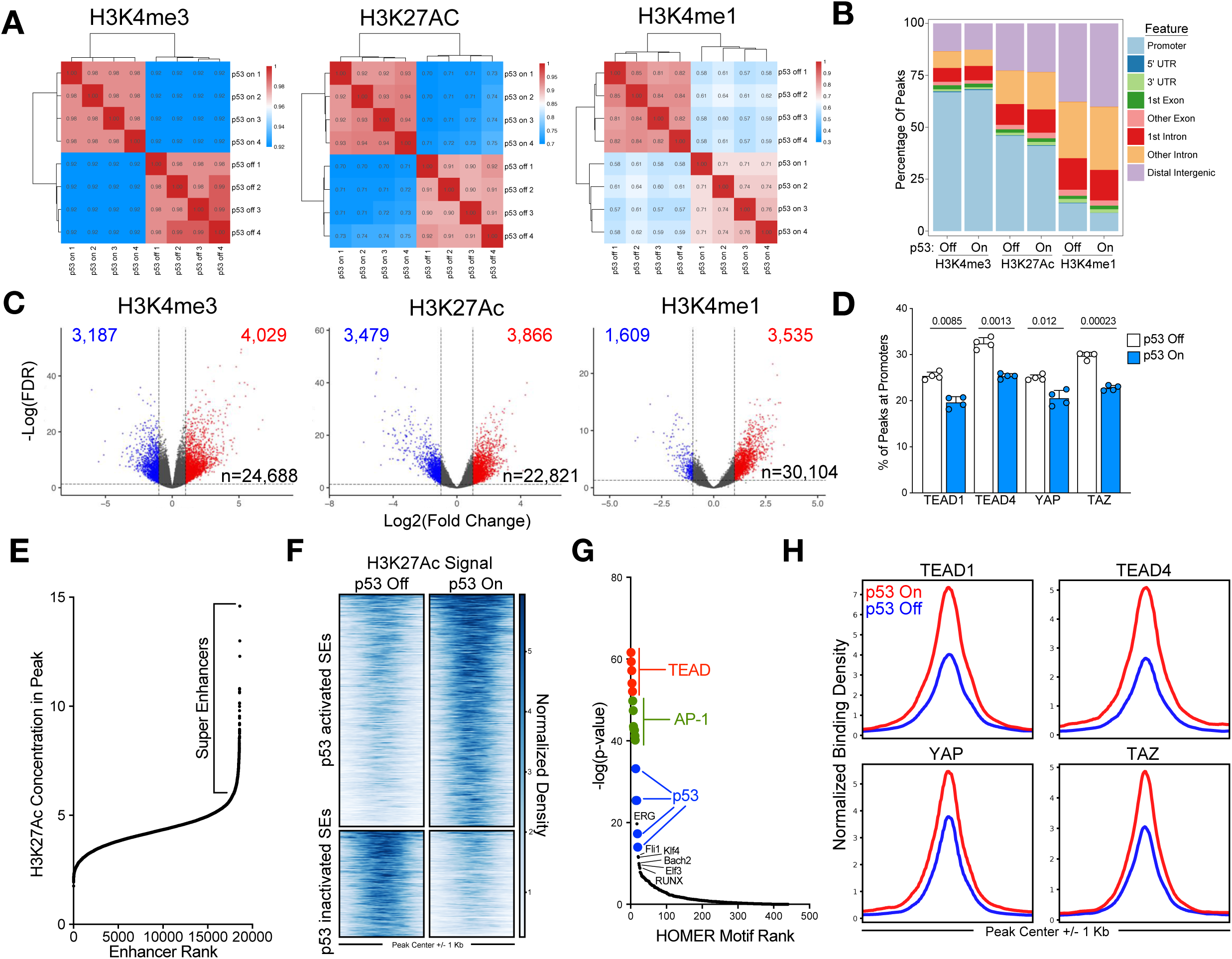
p53 restoration triggers chromatin remodeling. **(A-C)** CUT&RUN for H3K4me3, H3K27Ac, and H3K4me1 on dox (p53 off) and 6 days off dox (p53 on) in KP^sh^1 cells. **(A)** Heatmaps showing correlation of H3 mark peak densities across CUT&RUN replicates from four plates. **(B)** Plot depicting genomic annotation of consensus peaks. **(C)** Volcano plots of differential H3 mark peak densities upon p53 restoration with dots colored in red and blue denoting peaks with significantly increased and decreased binding (FC>1.5, FDR<0.05, EdgeR). **(D)** Percentage of CUT&RUN peaks within 1 Kb of TSS (promoters) of each CUT&RUN replicate. **(E)** Graph of DiffBind calculated peak concentrations of H3K27Ac CUT&RUN demonstrating super enhancer calls in cells with p53 on. **(F)** Heatmaps showing normalized H3K27Ac signal at super enhancers with significantly altered H3K27Ac (FC>1.5, FDR<0.05) by p53. **(G)** Homer known motif analysis of p53 activated super enhancers. **(H)** Profile plots of normalized binding density of TEAD1, TEAD4, YAP, and TAZ CUT&RUN at p53 activated super enhancers.

**Supplementary Figure 4:**
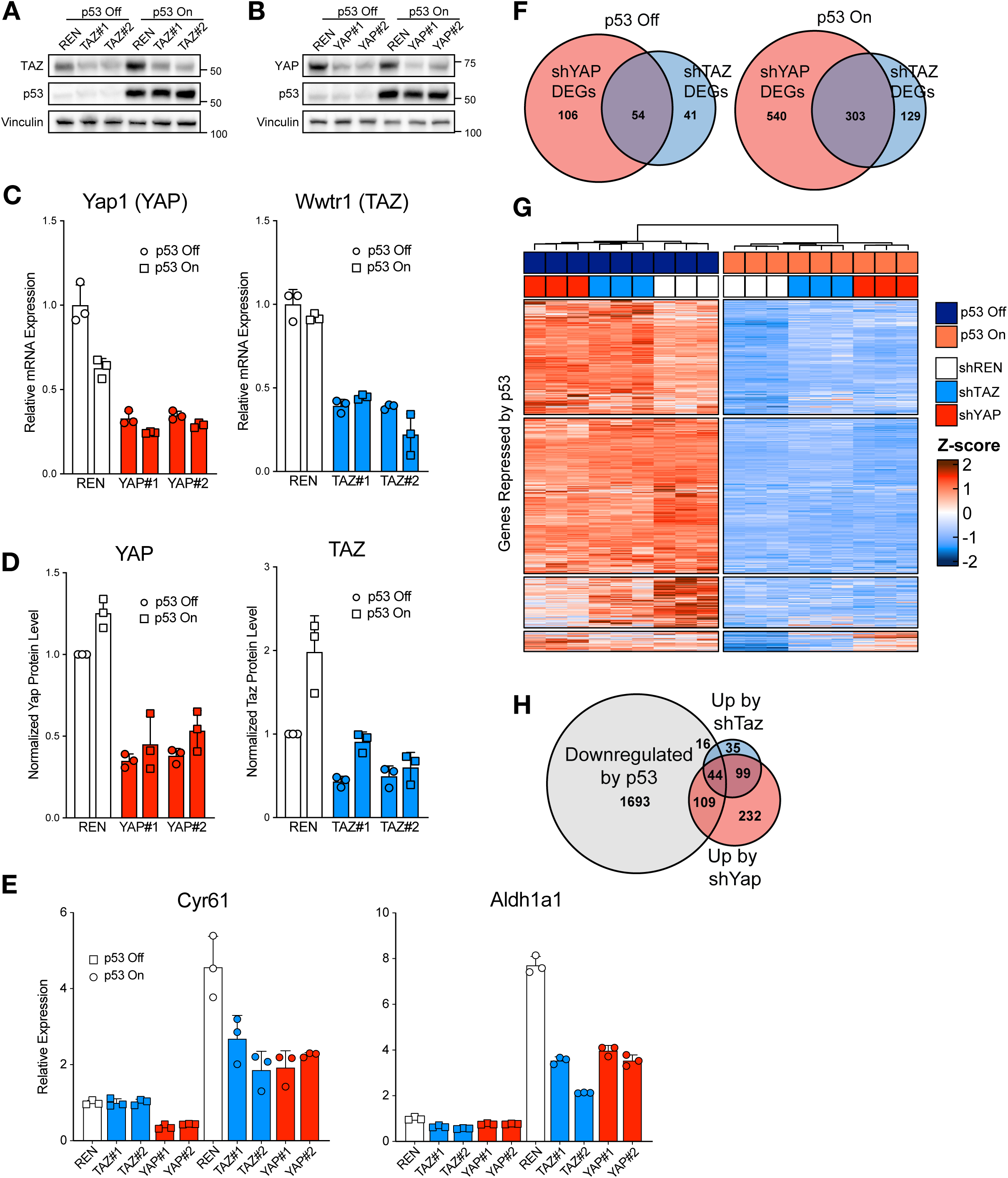
YAP/TAZ activity contributes to p53 gene regulatory networks. **(A-B)** Western blots of KP^sh^1 constitutively expressing shRNAs targeting **(A)** Wwtr1 (TAZ), or **(B)** Yap1 (YAP, right) vs a control shRNA against Renilla luciferase (REN) on dox (p53 off) or 6 days off dox (p53 on). **(C)** RT-qPCRs showing degree of TAZ and YAP mRNA depletion with shRNAs on dox (p53 off) and 6 days off dox (p53 on) from triplicate wells. **(D)** Quantification of Western blot band intensities showing degree of TAZ and YAP protein depletion with shRNAs on dox (p53 off) and 6 days off dox (p53 on) from three independent experiments normalized to beta Tubulin. **(E)** RT-qPCRs of TEAD target genes on dox or 6 days off dox (p53 on) in shRNA expressing cells normalized to Gusb. **(F)** Venn diagrams showing overlap of differentially expressed genes between shTAZ and shYAP with p53 off (left) and p53 on (right) (DESeq2, FC>1.5, FDR<0.05). **(G)** K-means clustered heatmap of all genes downregulated by p53 in shRenilla control cells (DESeq2 FC<-1.5, FDR<0.05). **(H)** Venn diagram of genes from (G) overlapping those with increased expression in shTAZ or shYAP cells with p53 on (DESeq2, FC>1.5, FDR<0.05).

**Supplementary Figure 5:**
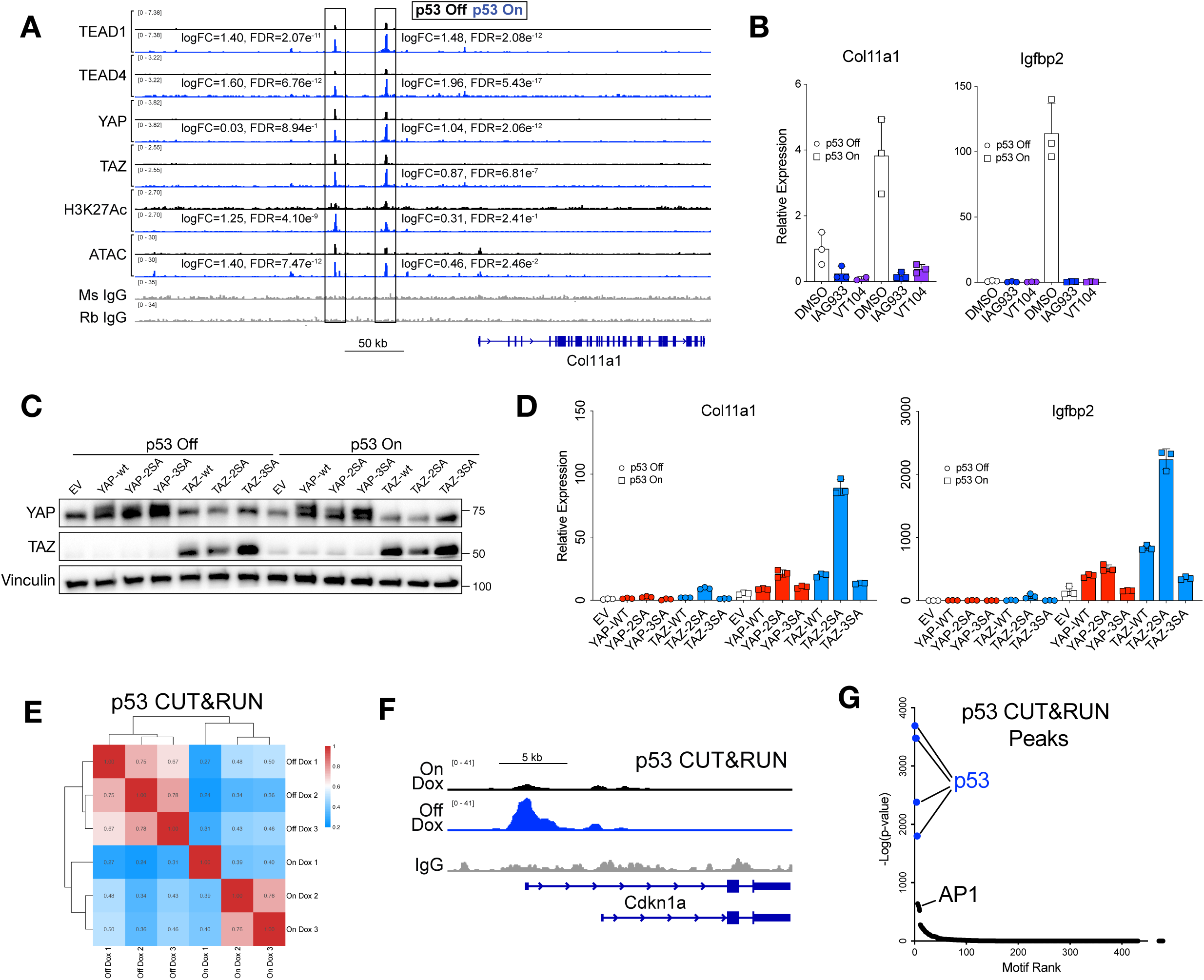
Co-regulation of genes by YAP/TAZ-TEAD and p53 binding. **(A)** Normalized averaged CUT&RUN and ATAC-seq signal in KP^sh^1 at the Col11a1 locus. **(B)** RT-qPCRs of KP^sh^2 treated with 62 nM IAG933 or 500 nM VT104 on dox (p53 off) or upon 6 days of dox withdrawal (p53 on). **(C-D)** KP^sh^2 were transduced with pBABE-Puro alone (EV) or containing various YAP/TAZ constructs. **(C)** Western blot of YAP/TAZ overexpression and **(D)** RT-qPCRs on dox (p53 off) and upon 6 days of dox withdrawal (p53 on) normalized to Gusb. **(E-G)** CUT&RUN for p53 on dox (p53 off) and 6 days off dox (p53 on) in KP^sh^1 cells. **(E)** Heatmaps showing correlation of p53 binding affinity across CUT&RUN replicates from triplicate wells. **(F)** Normalized average tracks of p53 binding at known target gene Cdkn1a with IgG control. **(G)** Homer known motif analysis of the p53 consensus peak list.

**Supplementary Figure 6:**
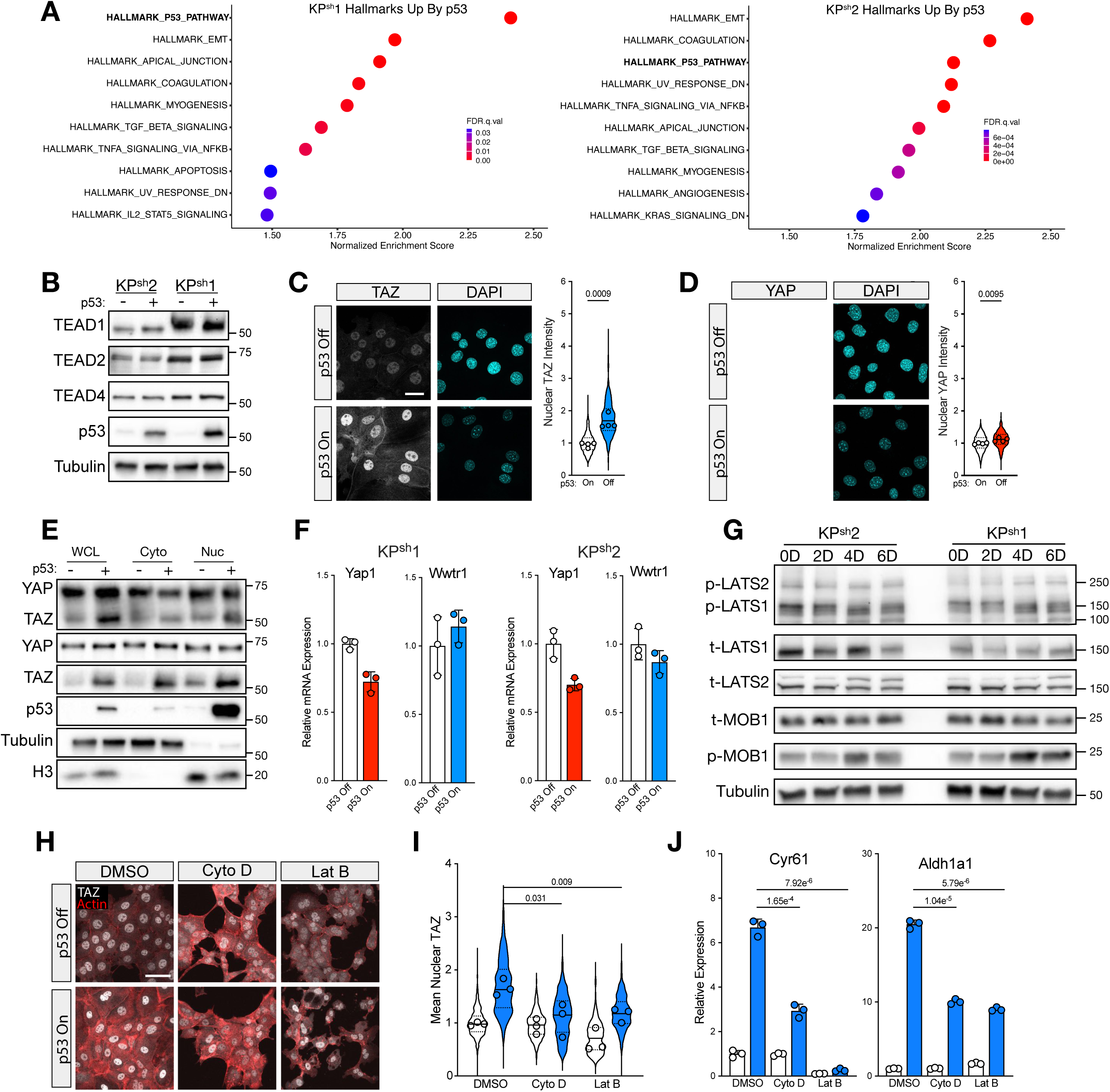
p53 restoration results in accumulation of nuclear TAZ. **(A)** Hallmark gene set enrichment analysis of RNA-seq from parental KP^sh^1 and KP^sh^2 cells upon 6 days of dox withdrawal for p53 restoration. **(B)** Representative western blots of TEAD levels in KP^sh^1 and KP^sh^2 cells on dox (p53 -) and 6 days off dox (p53 +). **(C-D)** IF microscopy images of KP^sh^2 subjected to 6 days of dox withdrawal (p53 on) and quantification of nuclear **(C)** TAZ and **(D)** YAP staining intensity. Dots show median signal across separate well of independent experiments and violin shows distribution combined across all experiments. p-value obtained using student’s t-test comparing median nuclear signal from quadruplicate experiments. **(E)** Western blots on whole cell lysate (WCL), Cytoplasm (Cyto), and Nuclear (Nuc) KP^sh^2 lysate cell fractions on dox (p53 -) and 6 days off dox (p53 +). **(F)** RT-qPCR of KP^sh^1 and KP^sh^2 for YAP1 (YAP) and Wwtr1 (TAZ) upon 6 days of dox withdrawal (p53 on) normalized to Gusb. **(G)** Representative western blots of Hippo pathway in KP^sh^1 and KP^sh^2 upon 6 days of dox withdrawal (p53 on). pLATS1-Ser909, pLATS2-Ser872, and pMOB1-Thr35. **(H-J)** KP^sh^2 treated with 12 hours of actin polymerization inhibitors Latrunculin B (Lat B, 5 µM) or Cytochalasin D (Cyto D, 1 µM) compared to vehicle (1:1000 DMSO) on dox (p53 off) or after 6 days of dox withdrawal (p53 on). **(H)** IF microscopy images and **(I)** quantification of IF microscopy images with dots showing median signal across separate wells of independent experiments and violin showing distribution combined across all experiments. p-value obtained using student’s t-test median nuclear signal from three independent experiments. **(J)** RT-qPCR normalized to Gusb, p-value from student’s t-test. Scale bars = 25 µm in (C,D) and 50 µm in (H).

**Supplementary Figure 7.**
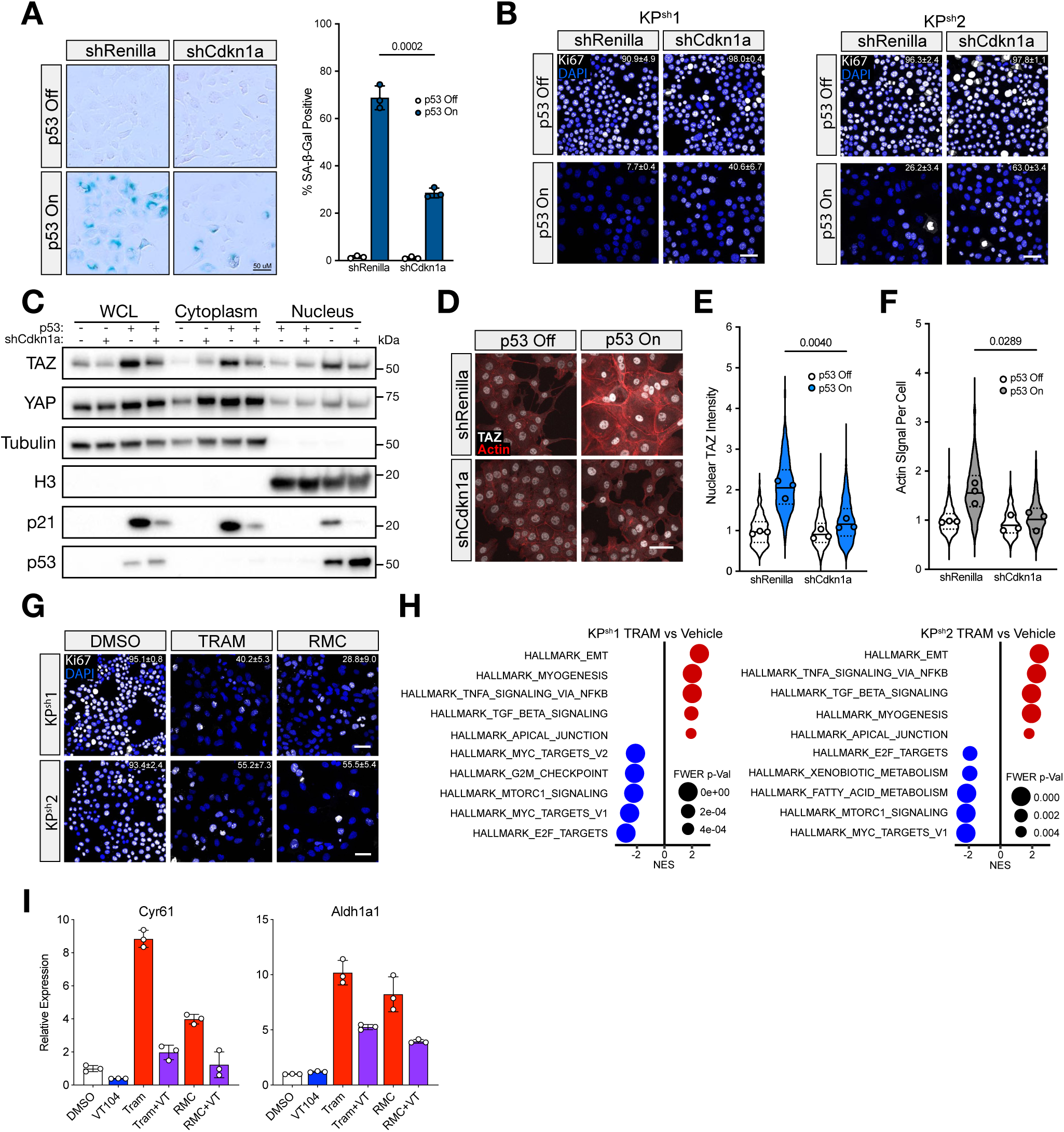
Senescence-induced actin reorganization facilitates TAZ activation. **(A-F)** KP^sh^ cells expressing shRNA targeting Renilla luciferase (control) or Cdkn1a. **(A)** Representative SA-β-Gal images of KP^sh^2 and quantification of three independent wells. p-value from student’s t-test. **(B)** Representative Ki67 IF images on dox (p53 off) and after 6 days of dox withdrawal (p53 on) with percentage Ki67 positive cells and standard deviation in upper right corner of image from quantification of three independent wells. **(C)** Western blot of whole cell lysate (WCL) or cell fractionation assay of cytoplasmic and nuclear fractions on dox (p53 -) and upon 6 days of dox withdrawal (p53 +) in KP^sh^1. **(D)** Representative IF microscopy images of TAZ and Phalloidin staining on dox (p53 off) and 6 days off dox (p53 on) with associated quantification of **(E)** nuclear TAZ and **(F)** total Actin per cell in KP^sh^2. Dots show median signal across separate three independent experiments and violins show distribution combined across all experiments. p-value obtained using student’s t-test of median signal from triplicate experiments. **(G)** Ki67 IF images of KP^sh^ treated with 72 hours of 50 nM Trametinib, 50 nM RMC, or DMSO vehicle control with percentage Ki67 positive cells and standard deviation in upper right corner of image from three wells. **(H)** GSEA showing the top five most enriched hallmark gene sets from RNA-seq in KP^sh^ treated with 25 nM trametinib or DMSO vehicle control for 72 hours. **(I)** RT-qPCRs from KP^sh^2 treated with 72 hours of 50 nM Trametinib, 50 nM RMC, or DMSO then an additional 48 hours with or without the addition of 500 nM VT104. Scale bars = 50 µm in (A,B,D, and G).

**Supplementary Figure 8:**
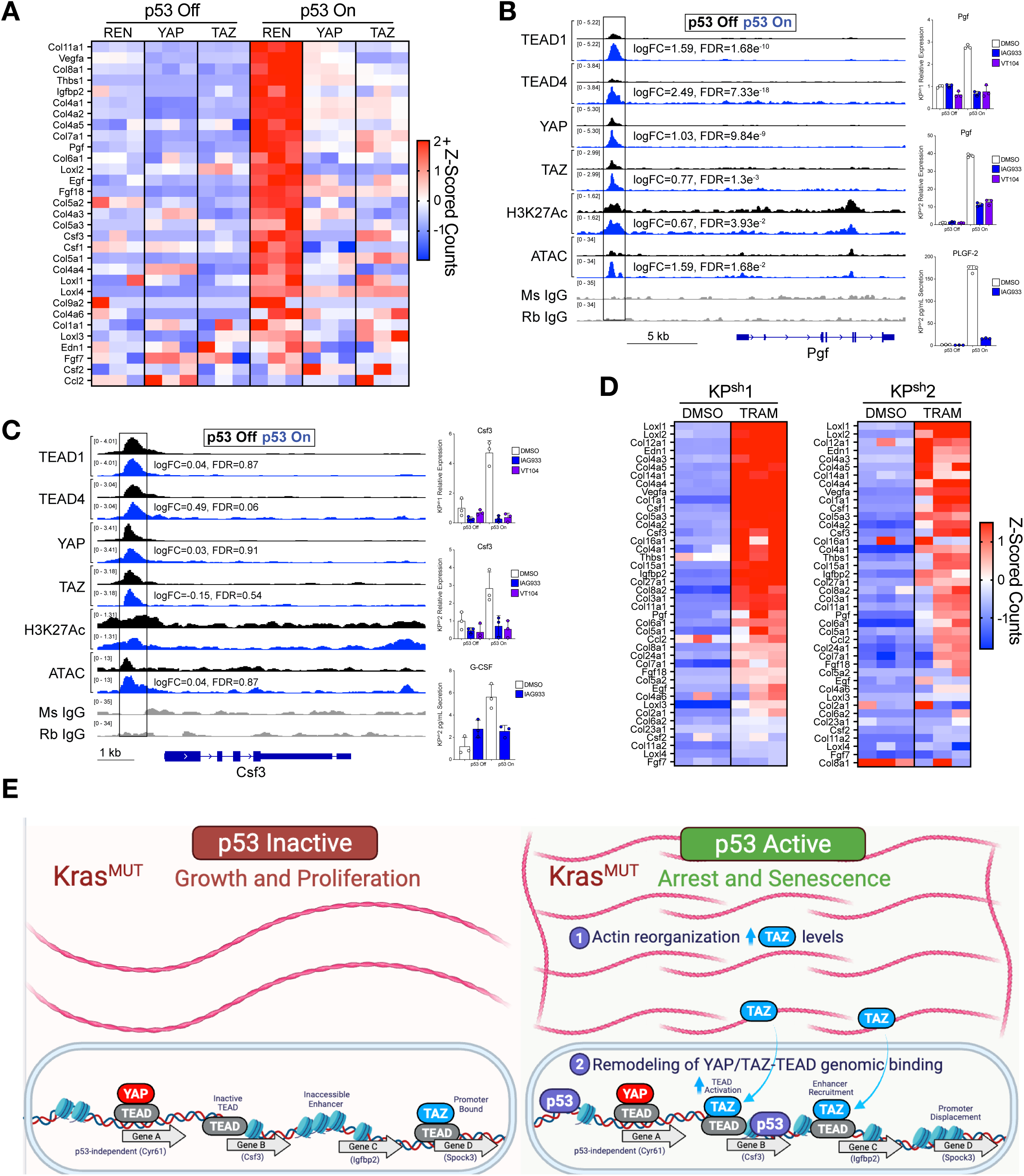
YAP/TAZ-TEAD regulates the SASP. **(A)** Heatmap of normalized Z-scored counts from RNA-seq of relevant secreted factors, collagens, and collagen cross-linking enzymes. **(B-C)** Normalized averaged CUT&RUN tracks (left), expression in KP^sh^1 (top right) and KP^sh^2 (middle right) from Figure 7E, and protein secretion in KP^sh^2 (bottom right) from Figure 7H for **(B)** Pgf/PLGF-2 and **(C)** Csf3/G-CSF. **(D)** Heatmap of normalized Z-scored counts from RNA-seq of KP^sh^ treated with 72 hours of 25 nM trametinib (TRAM) showing relevant secreted factors, collagens, and collagen cross-linking enzymes. **(E)** Model showing the two levels of p53-dependent control of YAP and TAZ.

## Methods

### Cell Culture

KP^sh^ cells were previously generated^15^ and passaged on dishes coated with collagen (PureCol, Advanced Biomatrix, 0.1 mg/mL) in complete DMEM containing 10% tetracycline-free fetal bovine serum (FBS, Gibco) and 1x penicillin-streptomycin. KP^sh^ were maintained on 1 ug/mL doxycycline (dox) until p53 restoration experiments whereby dox was removed for 6 days unless stated otherwise.

### Retrovirus and Lentivirus Production for Delivery of shRNAs and cDNAs

HEK293Ts were transfected using Lipofectamine 3000 (Thermo Scientific) according to manufacturer’s protocol for co-delivery of cDNA or shRNA cargo alongside packaging plasmids for virus production. Retrovirus was generated using pBS-CMV-GagPol (Addgene #35614) and pCMV-VSVG (Addgene #8454) while lentivirus was generated using psPAX2 (Addgene #12260) and pMD2.G (Addgene #12259). Guide-strand sequences of shRNAs targeting Renilla luciferase, Yap1, Wwtr1, and Cdkn1a/p21 (Supplementary Table 1) were cloned into the retroviral LMNe-BFP backbone^15^. Retrovirus was placed on KP^sh^ cells for 48 hours in the presence of 16 ug/mL polybrene and selected with 1 mg/mL geneticin/G418 for at least 4 days. Wildtype and phospho-null YAP and TAZ cDNAs were a gift from Channing Der^23^. YAP/TAZ cDNAs were cloned into pBABE-puro then retrovirus was produced for infection of the KP^sh^ cells in the presence of polybrene followed by selection with 10 ug/mL puromycin for 7 days.

### Western Blotting

Cells were washed with cold PBS then scraped into RIPA buffer (Thermo Scientific) containing cOmplete Protease Inhibitor Cocktail (Roche) and PhosSTOP (Sigma-Aldrich). BCA assay was performed to equalize protein concentrations across lysates before addition of β-mercaptoethanol containing Laemmli buffer (BioRad). SDS-PAGE was performed to separate proteins followed by wet transfers onto 0.45 uM PVDF membranes (Immobilon P). Membranes were blocked with a 5% solution of either fat-free milk or bovine serum albumin (phospho antibodies) rocking for a minimum of one hour at room temperature. Primary antibodies (Supplementary Table 2) were incubated in blocking solution overnight at 4 degrees C. After washing in TBST, secondary antibodies were added at 1:10,000 in blocking solution rocking for 1 hour at room temperature. Blots were then washed with TBST and imaged using a chemidoc (BioRad) after incubation in Clarity ECL reagent (BioRad).

### Growth Curves

KP^sh^ cells were removed from dox for 2 days to wash out the Trp53-targetting shRNA then 5,000 on dox and 15,000 off dox cells were seeded into 24-well plates. Cells were trypsinized and triplicate wells were counted using a Guava easyCyte at indicated timepoints. Fold change cell number is relative to the first cell count a day after seeding.

### Nuclear Cytoplasm Fractionation

Cells were subjected to the NE-PER nuclear and Cytoplasmic Extraction Reagent Kit (Thermo Fisher) according to manufacturer’s protocol with slight modifications. After isolating the cytoplasmic fraction, nuclei were resuspended in 250 mM sucrose, 10 mM MgCl_2_ containing 1x protease inhibitor tablet (Roche) and gently floated on top of 350 mM sucrose, 0.5 mM MgCl_2_ before centrifugation for 10 minutes at 1000 xg to remove contaminating cytoplasmic debris. Cold PBS was used as an additional nuclei wash followed by resuspension in Nuclear Extraction Reagent (NES). Sonication was used to shear nuclear DNA and insoluble particulate was cleared via centrifugation. Western blots were performed on fractions alongside whole cell lysates prepared as described above.

### Immunofluorescence Microscopy

KP^sh^ cells were seeded into ibi-Treated 4 well µ-slides (Ibidi) coated with a PureCol dilution of 1:30 in water. Fixation was performed with fresh 4% paraformaldehyde in PBS before washing with PBS and permeabilizing membranes with 0.3% Triton-X100 in PBS. Cells were blocked in 5% bovine serum albumin (BSA) dissolved in PBST. Antibodies were diluted in 3% BSA in PBST and incubated overnight at 4 degrees C. Primary dilutions were as follows: 1:250 TAZ (E9J5A) (CST, 72804) and 1:350 YAP (D8H1X) (CST, 14074). Secondary antibodies were incubated for 1 hour at room temperature protected from light in 5% BSA in PBST at 1:500 dilution. Additionally, a 10-minute incubation was performed with 1:2500 Phalloidin-iFluor 594 Reagent to stain actin (Abcam, ab176757) and either 1 ug/mL DAPI in PBS (Sigma, D9542) or 1:30,000 SYTOX green in TBS (Thermo Fisher, S7020) to visualize the nucleus. Cells were imaged using an Olympus IX81 or FV1000 Confocal Microscope (Olympus). Quantification of images was performed with Fiji (ImageJ). Nuclear YAP or TAZ levels were determined by measuring signal in the DAPI positive areas (nuclei), excluding anuclear mitotic DNA. Actin levels were determined by manually encircling individual cells and measuring average actin intensity across the cell body.

### Senescence Associated beta Galactosidase (SA-β-Gal) Assay

Cells seeded into 6-well tissue culture plates were washed with PBS then fixed with 0.5% glutaraldehyde in PBS at 37 degrees C for 15 minutes. Acidic PBS (pH=5.5, 1 mM MgCl_2_) was used to wash away the fixation solution before addition of X-Gal staining buffer containing 1 mg/mL X-gal dissolved in DMF, 5 mM potassium ferrocyanide, and 5 mM potassium ferricyanide diluted in acidic PBS. Staining was performed for 15 hours overnight at 37 degrees C before washing with PBS and imaging with an Olympus IX70 microscope. Percent SA-β-Gal cells were determined by counting stained cells manually with the Fiji (ImageJ) cell counter plug in.

### RT-qPCR

Cells were scraped into RLT lysis buffer containing beta mercaptoethanol, homogenized with QIAshredders (Qiagen), and RNA was extracted with RNeasy mini kit (Qiagen) according to manufacturer’s protocol. A NanoDrop One (Thermo Fisher) measured RNA concentrations followed by synthesis of 1 ug of cDNA with Reverse transcriptase (iScript, BioRad). RT-qPCR was performed on a CFX Opus Real-Time PCR System (Bio-Rad) with PowerUp SYBR green (Thermo Fisher). RT-qPCR primers were designed with MGH Harvard’s PrimerBank and can be found in Supplementary Table 1.

### RNA-seq of KP^sh^ cells

Three experiments were performed with the following conditions: parental KP^sh^1 and KP^sh^2 cells on dox or 6 days off dox, KP^sh^1 expressing shRNAs targeting Renilla, Yap1, or Wwtr1 on dox and 6 days off dox, and parental KP^sh^1 and KP^sh^2 cells treated with 72 hours of TRAM, vehicle on dox, or vehicle 6 days off dox. Samples were collected from triplicate wells by scraping into RLT lysis buffer containing beta mercaptoethanol, homogenized with a QIAshredder (Qiagen), and RNA was extracted with RNeasy mini kit. PolyA enrichment was followed by cDNA preparation and sequencing on a NovaSeq PE150. Raw data was processed using Partek Flow analysis software version 10.0.23.0720. Bases were trimmed for quality score > 20 and STAR version 2.7.8a was used to align to the mouse genome mm10. Gene counts were generated with HTseq followed by differential analysis by DESeq2 of all genes exhibiting the lowest maximum coverage of 2.0 counts. PCA plot was generated using transcripts per million of all differentially expression genes by ANOVA p<0.05. Gene Set Enrichment Analysis (version 4.3.2) was performed on median of ratios normalized counts using default parameters with 1000 gene set permutations.

### Autochthonous KPC^LOH^ Tumor model for RNA-seq

Mice containing p48-cre, LSL-Kras^G12D^, p53^flox^, CHC, and CAGs-LSL-RIK strains were maintained on mixed Bl6/129J as described previously^5^, fed dox chow (625 mg/kg, Harlan Laboratories), and subjected to small animal ultrasound until cancer formation. Pancreas tumors were finely chopped, rinsed with PBS, and dissociated into single cells rotating at 37 C for 45 minutes in 1 mg/mL Collagenase V (Sigma, C9263) and 1 mg/mL Dispase II (Sigma, D4693) dissolved in PBS containing 0.1 mg/mL DNase I (Sigma, DN25). Following a FACS buffer wash (10 mM EGTA, 2% FBS, 0.1 mg/mL DNAse I in PBS), cells were resuspended in 0.05% Trypsin-EDTA (Gibco) for 5 minutes at 37 C. Cells were then washed with FACS buffer, filtered through a 0.40 um membrane, and resuspended in FACS collection buffer (10 mM EGTA, 2% FBS, 0.1 mg/mL DNAse I in DMEM), with 1 ug/mL DAPI for dead cell exclusion. Live single positive GFP-/mKate2+ p53 LOH (SP) cancer cells and double positive GFP+/mKate2+ p53 intact (DP) cells from the same mice were sorted using a FACS Aria III (UNC flow cytometry core facility) using the gating strategy as described previously^5^. Pelleted cells were resuspended in RLT buffer with BME, homogenized with QIAshredders (QIAgen, 79656), and stored at −80C until RNA isolation using All prep RNA/DNA Micro Kits (QIAgen, 80284). Library preparation and RNA-seq was performed with Novogene’s HiSeq PE150 platform.

### KPC^LOH^ RNA-seq analysis

Raw sequencing data was processed using Partek Flow 12.7.0. Bases were trimmed using a quality score >20 and minimum read length of 25 bp. Alignment with STAR 2.7.8a was followed by counting with HTSeq 0.11.0. R was then used for performing within-sample normalization on raw counts such that the upper quartile of normalized gene expression counts was fixed at 1000 for each sample, followed by log2(x+1) transformation, to produce the log normalized counts. The PCA plot was generated by applying PCA to the log normalized counts from the top 2,000 features with the most variance. Differential expression analysis was performed on RNA-seq count data (unnormalized, unlogged) using *edgeR*. Matched DP-SP samples from the same mouse were paired, so the mouse sample ID was modeled as a blocking variable in the design matrix to account for the different baseline gene expression levels between mice. We fitted a negative binomial generalized linear model (NB-GLM), in which the response variable was the count data for each gene and the predictor variables were the sample ID and the p53 condition (SP p53 LOH vs DP p53 wild type). *EstimateDisp* and *glmQLFit* functions were used to fit NB-GLM, and the *glmQLFTest* function was used to perform differential expression testing by the quasi-likelihood-ratio test^67,68^. For GSEA, a pre-ranked gene list was generated by using the log(FDR) from the differential expression analysis with the sign of the fold change (signatures listed in Supplementary Table 3).

### CUT&RUN-seq

CUT&RUN was performed using the Cell Signaling Technology CUT&RUN assay kit according to manufacturer’s protocols. Briefly, KP^sh^1 cells were lightly fixed with 0.1% paraformaldehyde for 1 minute followed by quenching with 125 mM glycine and washing with PBS. Cells were scraped, and 500,000 cells were bound to Concanavalin A coated beads before incubation in primary antibody overnight at 4C. Secondary antibodies were added for 1 hour at 4C, bound by pAG-MNase for 1 hour at 4C, then DNA was digested for precisely 30 minutes at 4C by additional of CaCl_2_. Digestion was halted with STOP buffer and chromatin fragments were released for 30 minutes at 37 degrees. DNA crosslinking was reversed by addition of SDS and Proteinase K for 3 hours to overnight at 65 degrees C followed by purification using the ChIP DNA Clean & Concentrator kit (ZYMO). The Kapa Hyperprep Kit (Roche) was used for library preparation using half volumes as recommended by manufacturer’s protocol. A Qubit (Invitrogen) was used to determine DNA concentrations and a TapeStation (Agilent) validated fragment base pair size and integrity. Libraries were pooled and sequenced on a Nextseq 500 (p53) or Nextseq 1000 (YAP, TAZ, TEAD1, TEAD4, H3K4me1, H3K4me3, H3K27Ac) with a 75-cycle high output kit v2.5 or NextSeq 1000/2000 P2 reagents 100 cycle v3 kit respectively using paired sequencing. CUT&RUN was performed from triplicate (p53) or quadruplicate (YAP, TAZ, TEAD1, TEAD4, H3K4me1, H3K4me3, H3K27Ac) biological plates to confirm reproducibility.

### CUT&RUN data analysis

Raw sequencing data was demultiplexed to generate fastq files that were trimmed with TrimGalore (0.6.7) using -j –fastqc –paired –gzip –basename. Alignment to the mm10 genome was performed with bowtie 2 (2.3.4.1) –very-sensitive-local -X 800. Samtools aligned and sorted BAM files and the reads with mapq scores < 10 were removed. The csaw package was used to normalize independent samples using TMM on high-abundance regions (peaks) through the generation of scale factors flagging --peaks –winSize=200 --maxFrag=500 --pe=both. These scale factors representing effective library size within high abundance peak regions were divided by a million and the reciprocal was supplied to –scaleFactor in csaw to generate bigwig files. Replicate bigwigs were averaged with WiggleTools (n=3 for p53 or n=4 for all others per condition) and used for visualization via sample correlation heatmaps (DiffBind), peak density heatmaps (deeptools), profile plots (deeptools), or tracks (IGV). Peak calling was performed with MACS2 (2.2.7.1) with an FDR of 0.05 and blacklisted peaks were removed. Diffbind was used for differential occupancy or abundance analysis of transcription factors or histone marks respectively, removing grey listed peaks from the IgG control. For YAP, TAZ, p53, TEAD1, and TEAD4 CUT&RUN, a consensus peak set was generated by recentering individual peak files from all samples over a single summit with a 250 bp radius. These consensus summits were used for summit distance calculations. For H3K4me1, H3K4me3, and H3K27Ac CUT&RUN, a consensus peak set was generated by setting summits=False to maintain peak widths representative of wider enrichment of histone modifications. Peaks were only included if the peak was detected in at least 3 replicates for YAP, TAZ, TEAD1, TEAD4, H3K4me1, H3K4me3, and H3K27Ac or in at least 2 replicates for p53. Differential binding analysis on the consensus peak sets was performed with normalize=DBA_NORM_NATIVE and background=FALSE using the default EdgeR methods to generate a log2 fold change binding/abundance and adjusted p-value (FDR) upon restoring p53. ChIPSeeker was used to for peak annotation. Peaks annotated to genes for predicted regulation were filtered to −10 kb upstream through the 3’ UTR. Peaks were annotated to promoters if they overlapped a H3K4me3 peak within 1000 kb from a TSS and enhancers if they overlapped a H3K4me1 peak and were greater than 1000 kb from the nearest TSS. Correlation plots were generated via DiffBind’s plot function for peak affinity. Differential shared binding of TEAD1, TEAD4, YAP, and TAZ upon p53 restoration was identified by overlapping peaks with a 1.5-fold change and adjusted p-value < 0.05 identified in all four proteins. For motif enrichment analysis, homer known motif analysis was performed and reported by ranking the motifs based on FDR. Super enhancers were defined as non-promoter H3K27Ac peaks with H3K27Ac signal exceeding the rank-ordered geometric mean concentration counted and normalized by the DiffBind package in R. The threshold of p53 activated and inactivated SEs were set at FC>1.5, FDR<0.05 from SEs defined from the p53 on condition and FC<-1.5, FDR<0.05 defined from the p53 off condition.

### ATAC-seq

ATAC-seq data were processed using the ENCODE ATAC-seq pipeline^69^ implemented in the Workflow Description Language and executed with Caper^70^. Paired-end FASTQ files from (Morris 2019, GSE114263) were subjected to the standardized workflow including adapter trimming, alignment with Bowtie2 to mm10^71^, and duplicate removal. MACS2 was utilized for peak calling of ATAC-seq at an FDR of 0.05. Bedtools was used to generate a consensus accessibility peak list containing overlapping peaks found in at least 2 replicates. DiffBind was used for differential accessibility analysis by centering peak summits with a 250 bp radius from the consensus peak list. The default EdgeR method was used for normalization and differential binding analysis flagging normalize=DBA_NORM_NATIVE and background=FALSE. Loci with increased accessibility > 1.5-fold and adjusted p-value < 0.05 were used for motif enrichment analysis using HOMER known motif profiling. Profile plots were generated with deeptools by generating a matrix of accessibility over the shared differentially bound TEAD1/TEAD4/YAP/TAZ peak list.

### Analysis of ATAC-seq from pancreatic Kras mutant pre-malignant and malignant cells

Normalized ATAC-seq peak counts were obtained from the NCBI accession number GSE132440^13^. The sorted pancreatic epithelial cells (mKate2+, GFP+) from pre-malignant p53 intact samples were comprised of KC;RIK mice treated with 2 days of caerulein (n=4) and the PDAC samples were generated from cancer arising in KP^fl^C mice (Ptf1a-cre;RIK;LSL-KrasG12D;p53fl/+) (n=1) or after syngeneic orthotopic transplantation of Kras^G12D^;p53^KO^ engineered pancreatic organoids (n=3). Differential accessibility analysis was performed with DiffBind between pre-malignant and malignant peak sets. Homer known motif analysis was used to identify loci with increased accessibility in pre-malignant samples compared to cancer (FC>1.5, FDR<0.05).

### Analysis of scRNA-seq of neoplastic and malignant pancreas tumors

scRNA-seq data generated by Burdziak et al. 2023 was downloaded from Gene Expression Omnibus (GSE207943)^18^. We analyzed the entire spectrum of Kras-mutant epithelial cells from stages K1-K6. There are 13,650 p53-wild type K1-K4 pre-malignant cells and 8,855 p53 inactivated malignant cells from K5-K6. The standard *Seurat* pipeline^72^ was used to analyze the scRNA-seq data. The reported UMI counts were normalized for library size and log-transformed, and the log-normalized data from the top 5,000 variable genes were centered, scaled, and PCA was performed. To visualize scRNA-seq data, uniform manifold approximation and projection (UMAP) was performed on the top 50 principal components. The *AddModuleScore* function from the R package *Seurat* was used to compute expression scores of each gene signature for individual cells, and gene expression scores were then scaled from −0.5 to 1 across all cells. UMAPs were generated by projecting the gene expression scores or scaled individual gene expression score across the population of cells. Pearson correlation coefficients and significance was determined for the reported pairs of gene signatures.

### Cytokine Array

KP^sh^2 cells were seeded into 1:1000 DMSO or 125 nM IAG933 on dox or after 5 days of dox withdrawal. After 24 hours, 2 mLs of fresh complete DMEM containing drug was added for 48 hours for conditioning before collection of conditioned medium (CM) from triplicate wells. The cells were counted with a Guava easyCyte and CM was diluted to an equivalent cell number per medium volume. CM was analyzed using Eve Technologies’ Mouse Cytokine/Chemokine 32-Plex Discovery Assay® Array (MD32) and Mouse Angiogenesis and Growth Factor 16-Plex Discovery Assay® Array (MDAG16) (Factors analyzed, Supplementary Table 4).

## Notes

### Competing Interest Statement

The authors have declared no competing interest.

